# Rhizobial motility preference in root colonization of *Medicago truncatula*

**DOI:** 10.1101/2025.10.12.681932

**Authors:** Anaïs Delers, Anne Bennion, Ambre Guillory, Lisa Frances, Elizaveta Krol, Fanny Bonnafous, Laurena Medioni, Javier Serrania, Rémi Peyraud, Joëlle Fournier, Fernanda de Carvalho-Niebel, Anke Becker

**Author notes:** **Authors for correspondence:** Fernanda de Carvalho-Niebel and Anke Becker. These authors contributed equally to this work (Anaïs Delers, Anne Bennion).

## Abstract

- Tunnel-like infection thread (IT) structures support root colonization by symbiotic nitrogen-fixing rhizobia bacteria in most legume species. These tip-grown structures are key to direct rhizobia from root hairs to developing nodules, where they are hosted to fix nitrogen. Rhizobia likely progress inside ITs by combining growth and motility, by modes not yet defined. Here, we tackled this question by combining mathematical modeling, live cell imaging, and bacterial mutant phenotyping in *Medicago truncatula*.
- Modeling the motion of fluorescently-labeled *Sinorhizobium meliloti* inside early root hair IT compartments estimated slow movement (2 to 6 µm/h), compatible with passive rather than active motility. Consistent with this model, flagella-less *fliF* and *fliF-fliRdel S. meliloti* mutants were impaired in active swimming motility *in vitro,* yet could colonize host roots and nodules *in planta*. In contrast, mutation in the rhizobactin 1021 siderophore *rhbE* biosynthesis gene affected both surface motility *in vitro,* and host root and nodule colonisation. This mutation also promoted the formation of branched ITs in root hairs, which ultimately resulted in impaired nodule development and infection.
- Our findings support the model estimation and suggest that *S. meliloti* prioritises flagella-independent surface translocation, partially by secreting rhizobactin 1021 surfactants to reach developing nodules in *M. truncatula*.

## Introduction

Symbiotic relationships with soil microorganisms can help plants access the nutrients they need for growth. Certain angiosperm species in the nitrogen-fixing clade evolved the ability to obtain nitrogen through symbiosis with bacteria, which they host intracellularly in specialized organs called root nodules (Huisman & Geurts, 2020). These interactions have been well-studied in legume plants, notably in model species such as *Medicago truncatula*, which hosts the rhizobia symbiont *Sinorhizobium meliloti*. Establishing this interaction requires precise molecular exchanges between partners (Krönauer & Radutoiu, 2021) before bacteria can colonize their host, which occurs in most cases via a tunnel-like apoplastic compartment called the infection thread (IT) (Gage, 2004).

ITs initiate in root hairs and proceed through well-defined stages (reviewed in de Carvalho-Niebel *et al*., 2024). Root hairs curl around Nod factor-producing rhizobia (Esseling *et al*., 2003) and enclose them in a radially-expanding infection chamber where they proliferate until polarized secretion creates the tip-growing IT tubular structure (Fournier *et al*., 2015). ITs are sequentially re-initiated in successive cell layers to guide rhizobia transcellularly from root hairs to the developing nodule primordium, where they are released, endocytosed, and differentiated into N-fixing bacteroids (Yang *et al*., 2022).

The successful formation and progression of ITs within plant cells depends on the plant’s specific perception or controlled degradation of rhizobial Nod factors or exopolysaccharide signals (Kawaharada *et al*., 2017; Malolepszy *et al*., 2018). The plant host also triggers a series of cellular events (reviewed in de Carvalho-Niebel *et al*., 2024) to create the optimal IT apoplastic environment for rhizobia colonization. Inside the IT space, rhizobia progress in a sparse, single-file arrangement, slightly behind the IT tip that extends in a cytoplasmic bridge connected with the migrating nucleus (Fournier *et al*., 2008; Guillory *et al*., 2024). It has been proposed that rhizobia progress in the IT environment by combining cell proliferation and collective movement, though the form of motility they actually use is unknown.

Rhizobia, like other bacteria, use active, flagella-dependent or passive flagella-independent motility to move on surfaces (Wadhwa & Berg, 2022). Rotation of the rigid motor-driven flagella filaments powers individual bacteria to swim in liquid media or multiple bacteria to swarm across solid surfaces. Bacteria can also use passive sliding movement on surfaces, driven by the outward pressure of dividing cells and compounds (e.g., surfactants) that help to reduce friction between cells and surfaces. In *S. meliloti*, active motility is enabled by 8 peritrichous flagella (Gotz & Schmitt, 1987). These are formed under the control of the flagellar regulon, which comprises flagellar, chemotaxis, and motility genes, grouped in one contiguous 45-kilobase chromosomal region and regulated in a three-class hierarchical manner (Sourjik *et al*., 1998, 2000). Motility in *S. meliloti* is also regulated by the ExpR/Sin quorum-sensing system, which can downregulate the expression of flagellar biosynthesis genes at high population density (Hoang *et al*., 2004). This quorum-sensing system also regulates the production of exopolysaccharides (Hoang *et al*., 2004), including symbiotically active EPS II (galactoglucan), which promotes swarming motility (Gao *et al*., 2012). *S. meliloti* strains 1021 and 2011, frequently employed for symbiotic studies in *M. truncatula*, are disrupted in *expR* (Pellock *et al*., 2002). Although EPSII biosynthesis is reduced in these strains, swarming motility is still observed, albeit dependent on the biosynthesis of the siderophore rhizobactin (Rhb) 1021 (Nogales *et al*., 2010, 2012). Thus, Rhb1021 may act as a surfactant to facilitate surface motility.

Siderophores are high-affinity iron chelators secreted by many organisms, including bacteria (Timofeeva *et al*., 2022). Rhb1021 is composed of a modified citrate backbone synthesized by enzymes encoded on the *rhbABCDEF* operon (Lynch *et al*., 2001). After scavenging iron, siderophores bind an outer membrane receptor before being pumped back into the cell (Timofeeva *et al*., 2022). In *S. meliloti*, Rhb1021 uptake depends on an outer membrane receptor encoded by *rhtA* and a permease encoded by *rhtX* (Lynch *et al*., 2001; Cuív *et al*., 2004). While mutations of Rhb1021 biosynthesis genes in ExpR-deficient *S. meliloti* strains abolish surface motility, it is not affected when only RhtA-mediated siderophore uptake is prevented, suggesting that the Rhb1021 function for surface motility lies outside the cell (Nogales *et al*., 2010).

Motility is also critical for rhizobia’s symbiotic interaction with their legume hosts. Flagella motility helps rhizobia in chemotactic movement to host roots, colonization, and attachment to root surfaces as well as increased competitiveness for nodule occupancy (Ames & Bergman, 1981; Mellor *et al*., 1987; Caetano-Anollés *et al*., 1988; Catlow *et al*., 1990; Fujishige *et al*., 2006; Miller *et al*., 2007; Zheng *et al*., 2015; Aroney *et al*., 2021, 2024; Compton & Scharf, 2021; Navarro-Gómez *et al*., 2024). Moreover, a transposon insertion sequencing genetic study in *Rhizobium leguminosarum* found that functional flagella genes enhance bacterial survival and growth in later stages of nodule development (Wheatley *et al*., 2020). But, so far, non-motile flagella mutants did not appear to significantly affect nodulation or nitrogen fixation, at least in alfalfa or clover species (Ames & Bergman, 1981; Mellor *et al*., 1987). Still, the role of flagella-dependent or -independent motility in rhizobia transcellular host infection remains unclear. Though Rhb1021 biosynthesis seems to enhance biofilm formation and the efficiency of nitrogen fixation in alfalfa (Gill *et al*., 1991; Amaya-Gómez *et al*., 2015), its impact during rhizobia plant host interaction remains largely unexplored.

In this study, we explored confocal time-lapse images of fluorescent *S. meliloti* infection events to infer speed and modes of motility used by rhizobia in transcellular IT compartments. To challenge this model, we generated a series of flagella or Rhb1021 biosynthesis and transport mutants in the ExpR-less *S. meliloti* strain 2011 to tackle the question of how flagella-dependent and independent paths impact early stages of nodule development and rhizobia colonization in *M. truncatula.* These strains, carrying constitutive β-galactosidase (*lacZ*) or fluorescent reporters, enabled the quantification of their ability to colonize root hairs or emerging nodules. Altogether, our findings provide novel insights into motility modes privileged by rhizobia to colonize their host roots.

## Materials and Methods

### Plant materials, bacterial strains and culture conditions

*M. truncatula* Jemalong A17 and the *super numeric nodules*-2 *sunn-2* mutant (Schnabel *et al*., 2005) were used in this work. The bacterial strains and plasmids used in this study are listed in Table **S1**. *Escherichia coli* strains were grown at 37°C in lysogeny broth (LB) medium (10 g/L tryptone, 5 g/L yeast extract, 5 g/L NaCl). *S. meliloti* strains were grown at 28-30°C in tryptone yeast (TY) medium (5 g/L tryptone, 3 g/L yeast extract, 0.4 g/L CaCl_­­_). When appropriate, the following antibiotics were added: kanamycin (25 mg/L for *E. coli,* 100 mg/L for *S. meliloti*), spectinomycin (50 mg/L for *E. coli,* 100 mg/L for *S. meliloti*), streptomycin (200 or 600 mg/L), tetracycline (10 mg/L).

### Generation of *S. meliloti* mutant strains

The constructs used in this work were generated using standard genetic techniques. Primers used for the genetic manipulations are listed in Table **S2,** and a detailed description of the plasmid construction is provided in Table **S3**. Plasmids were transferred to *S. meliloti* by conjugation with *E. coli* strains. S17-I was used for the conjugation of the pXLGD4 and pK18mobsacB-based plasmids, and DH5α with helper strain XL1-Blue were used to conjugate the pAB14 plasmid (described in Döhlemann *et al.,* 2016). Deletions of *fliF-fliR, rhbE*, and *rhtA* were generated using the pK18mobsacB suicide plasmid vector, which is non-replicative in *S. meliloti* and confers both sucrose sensitivity and kanamycin resistance (Schäfer *et al.,* 1994). For each deletion, DNA fragments upstream and downstream of the deletion target were inserted into pK18mobsacB (see Tables **S2** and **S3**). Double recombinant deletion strains were selected on LB agar supplemented with 10% sucrose, as previously described (Schäfer *et al.,* 1994). The mScarlet reporter was made by inserting the mScarlet coding sequence into the shuttle vector pABC-Psyn plasmid under a constitutive promoter (oligonucleotides 2 and 3, Table S2) from the 5’UTR (-50 to -1) of *SMc06412* and a leader sequence and ribosomal binding site from the 5’UTR of *mucR* (-44 to -1). Constructs were confirmed by PCR amplification and DNA sequencing.

### Genomic DNA isolation and sequencing

For genomic DNA (gDNA) isolation, *S. meliloti* cell cultures were grown overnight in TY with appropriate antibiotics until they reached an exponential phase, with an OD600 between 0.4 and 1. *S. meliloti* gDNA from (*Sm*2011-lacZ) WT and *fliF-fliRdel* mutant strains was extracted using a Blood & Cell Culture DNA Kit (Qiagen) with a modified protocol from (Mayjonade *et al*., 2016). gDNA from *rhbE* and *rhtA* mutants was purified using the NucleoSpin Microbial DNA Mini Kit (Macherey-Nagel). Finally, gDNA purity was verified by measuring gDNA concentration and spectrophotometric ratios (A260/A280; A260/A230). *Sm*2011-lacZ (WT) and *fliF-fliRdel* gDNA at 33ng/µL was then used for long-read sequencing on the Oxford Nanopore Technologies (ONT) PromethION (Table **S4**). *rhbE* and *rhtA* mutant gDNA at 50 ng/μL was sequenced by Plasmidsaurus using ONT (Table **S4**). Read mapping to the *Sm*2011 GMI11495 reference genome and comparisons for detection of sequence variants were carried out using CLC Genomic Workbench 24.0.1 (Tools: Map Long Reads to Reference 1.2, Basic Variant Detector 2.6 - applied thresholds: Coverage >= 10, Count >= 6, Frequency >= 75%).

### *S. meliloti in vitro* motility assays

Swimming motility assays were performed using cell cultures grown overnight in TY media supplemented with appropriate antibiotics. 2 µL of cell culture was spot inoculated on soft TY 0.3-0.4% agar plates (OD600nm = 1; for Fig. **3b**). Plates were imaged after incubating for 3-4 days at 28°C. Surface motility tests were performed on MM 0.6% on semisolid Noble agar plates (adapted from Bernabéu-Roda *et al*., 2021; Doin de moura, 2023), prepared as follows: 1.2% agar and 2X MM were autoclaved in a pressure cooker for 5 minutes at 55°C, then mixed and stirred for 5 min, followed by 15 min at 15°C, before adding the vitamins. Then 25 ml of medium was poured into each plate, and after drying for 10 minutes with the open plates, 2 µL of cell culture grown ON and then for 5 h in MM medium (OD600nm = 0.1) was spot inoculated and dried for 4 min. Univerted parafilmed plates were imaged after 72 h of incubation at 28 °C. The area (cm²) of each spot from two (swimming) or three (surface motility) technical replicates per experiment was measured using ImageJ software.

### Plant growth and bacterial inoculation

After pod dehulling, seeds were scarified by 95% sulfuric acid treatment for 10 min. After washing, bleach treatment (12% sodium hypochlorite) was performed for 2 min to facilitate seed sterilization. After rinsing several times, the seeds were kept in water for a few minutes before being sown on soft agar medium. The Petri dishes sealed with Parafilm were inverted for 2 to 6 days in the dark at 4°C to synchronize germination. Plates were then incubated at 20°C or 16°C for 17 h to 24 h to induce germination until radicles were more than 1 cm long, and then used for IT observation and nodulation experiments.

For IT observation using *in vivo* imaging, germinated seedlings were placed on Fahraeus medium plates (12 cm × 12 cm) supplemented with 0.5 mM NH_­­_NO_­­_, after their root tips were removed to promote new root emergence. Plates were placed vertically in plastic boxes with black plastic bags covering the roots and kept under controlled 16 h neon light/8 h dark photoperiod conditions and a light intensity of 70 mE/s/m2 at 20 °C. After 3 days to one week, plants with new root systems were transferred to nitrogen-free 0.5% [w/v] phytagel Fahraeus plates supplemented with 50 nM 2-aminoethoxyvinyl glycine (AVG), root systems were covered with a sterile LUMOX film (Sarstedt, UK), and the plates were placed with an inverted tilt so that the roots would grow along the film, as described previously (Fournier *et al*., 2015). After 3 days of nitrogen starvation, plant roots were inoculated with 800µL of the *Sm*2011-GFP (Fig. **1**) or the *Sm*2011-mScarlet WT or *rhbE* (Fig. **6**) cell suspension at OD600nm 0.001, directly onto the roots between the medium and the film. After 30 min of flat incubation, the inoculated plants were returned to the growth chamber with their root systems tilted downwards and kept in the dark until microscopic observations were made. Depending on the speed of root growth, plants were kept at 20°C until observation or moved to a 25°C chamber, 5 to 7 days after germination. During the observation period, the plants were placed in a phytotron under controlled 16 h neon light/8 h dark photoperiod at 25 °C and 35% humidity.

**Fig. 1.**
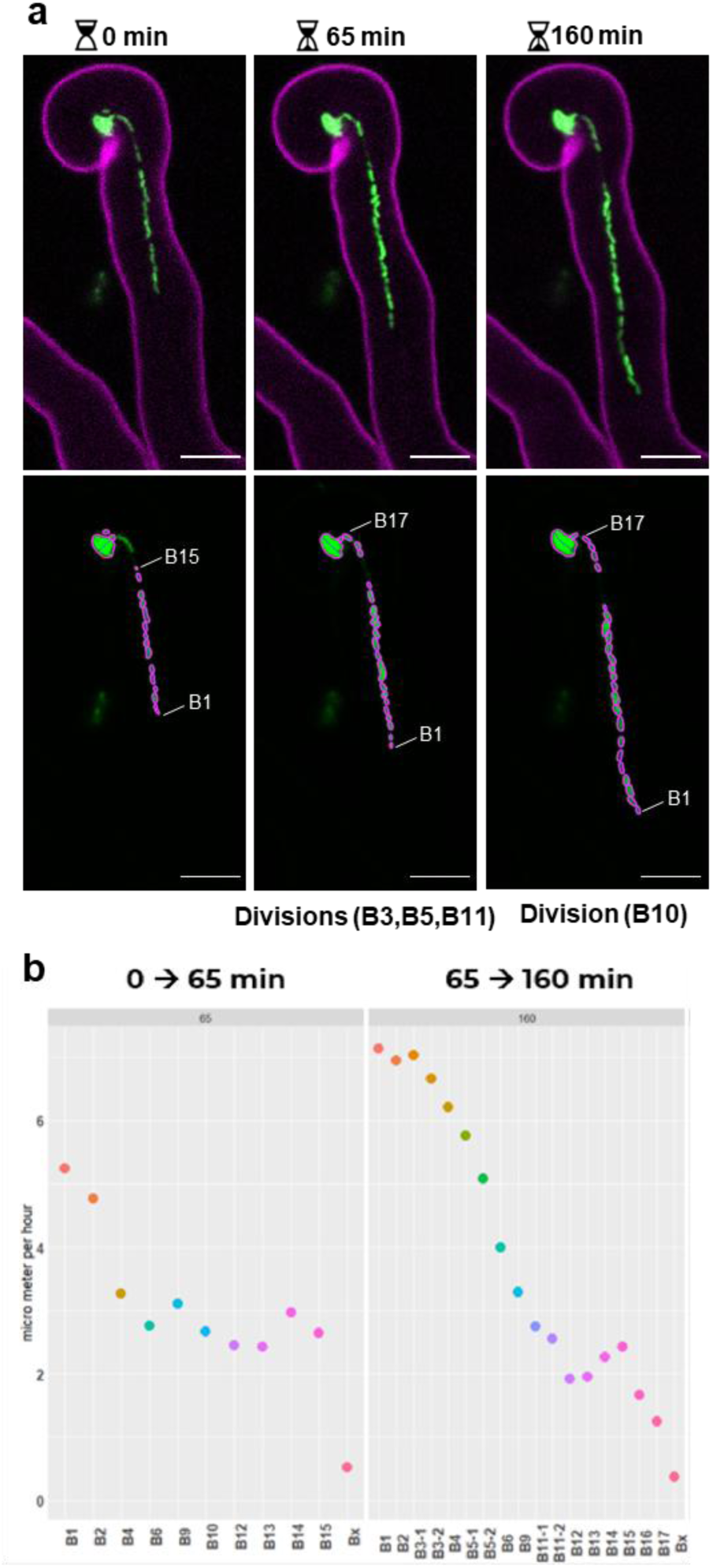
A model of rhizobia motion inside ITs. (**a**) Tracking bacteria within ITs. Upper panel: Three successive confocal images of a developing IT within an *M. truncatula* root hair were captured at 65 and 95 min intervals. Images of GFP fluorescence from the rhizobia strain (in green) and autofluorescence of the RH cell wall (in magenta) were merged, and maximal projections of 3 (2 first timepoints) or 6 (last timepoint) confocal sections are shown. Lower panel: The GFP channel of each IT image was used to define regions of interest (ROIs) corresponding to individual rhizobia within the developing IT. Rhizobia were subsequently numbered from the IT tip to the older part, close to the infection chamber (detailed in Suppl Fig. S1). The ROIs corresponding to the infection chamber and a neighbouring cell (Bx) were used as a spatial reference. Spatial coordinates of the bacterial cells were determined using the MicrobeJ plugin of ImageJ and used to calculate the speed of each cell over each time interval. Scale bars **(a)** = 10µm. See also Supplemental Table S8 and Fig. S1. (**b**) Assessment of the speed of bacterial cells inside ITs. The two graphs present the speed of some individual rhizobia and their offspring generated during the time frame of the experiment, between 0 and 65 minutes, left graph, and between 65 and 160 minutes, right graph. The speed of the individual cell was calculated using an agent-based mathematical model. See also Supplemental Table S1 and Fig. S1.

For nodulation experiments described in supplementary Figure **S2**, germinated seedlings of *M. truncatula* A17 were transferred to 8 × 8 × 7 cm pots (three plants per pot) filled with inert attapulgite substrate (Oil Dri US Special; http://www.oildri.com/) supplemented with 10 mL nitrogen-free Fahraeus medium then placed in small greenhouses at 25 °C, with a 16 h photoperiod and a light intensity of 100 mE/s/m2. After 3 days of nitrogen starvation, plants were inoculated with a suspension of *Sm*2011-lacZ WT or mutant strains (OD600nm 0.1; 4mL per pot). Plants were watered with sterile distilled water and harvested at 7 dpi.

For phenotyping experiments in Fig. **2**, **4**, **5** and **6**, germinated seedlings of *M. truncatula* A17 were transferred to 8 × 8 × 7 cm pots (three plants per pot) filled with a mixture of fine vermiculite substrate (∼2/3; Agrigaronne; https://www.agri-garonne.fr/) and sand (∼1/3; silica 0.7/1.3 mm; Puel) and then placed in trays hermetically covered with cellophane until the morning of inoculation in a chamber at 25°C day-22°C night, with a 16 h led light/8 h dark photoperiod, a light intensity of 320 µmol/m²/s and 55% humidity. After 3 days of nitrogen starvation, plants were inoculated with a suspension of *Sm*2011-lacZ WT or mutant strains (OD600nm 0.05; 20 mL per pot). Plants were watered twice a week with nitrogen-free Fahraeus medium (Figure **5**) or nitrogen-free Plant Prod (Fig. **3** and **S3**) (15%P-40%K; Agrigaronne; https://www.agri-garonne.fr/) and harvested at 5 or 7 dpi.

**Fig. 2.**
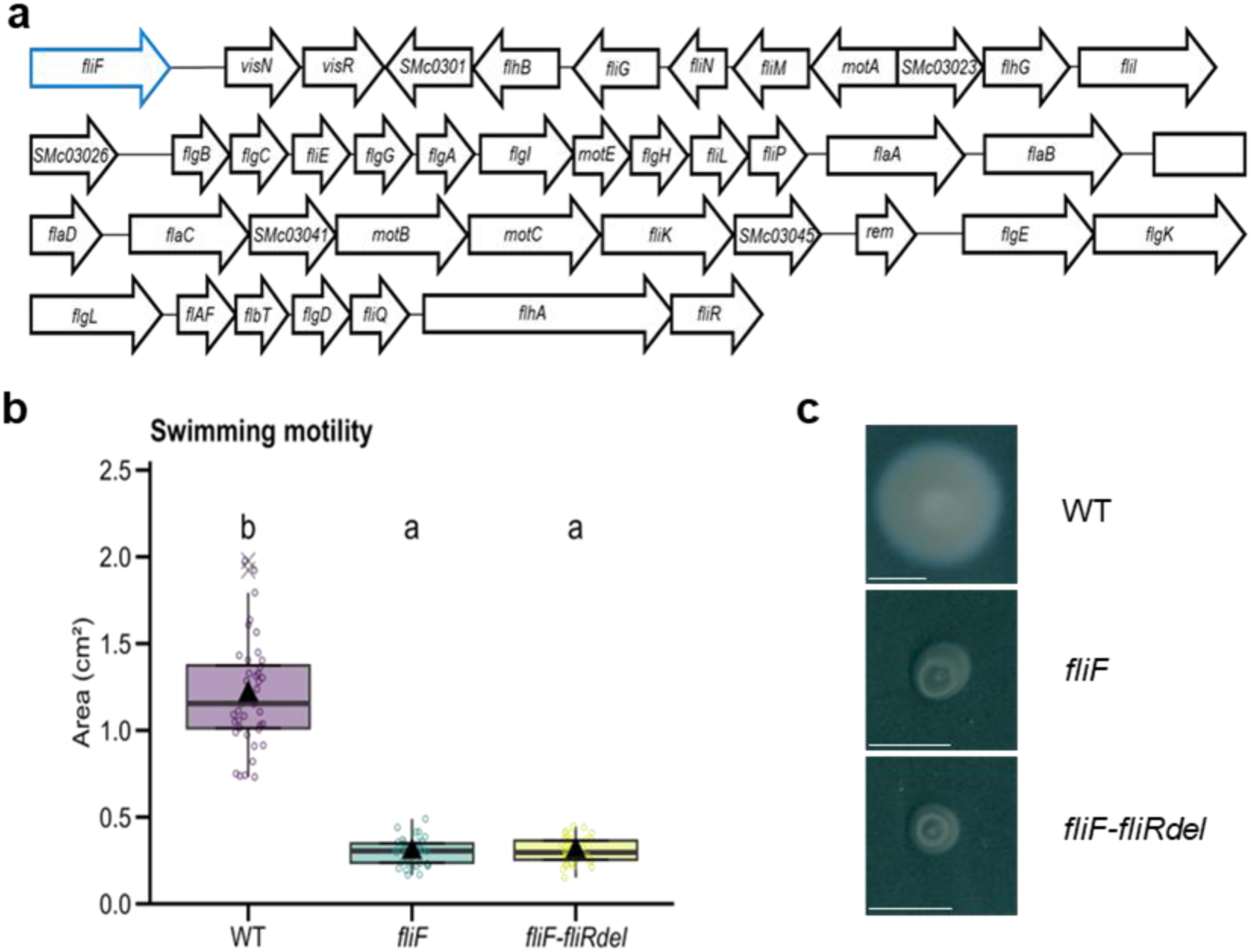
*S. meliloti fliF* and *fliF*-*fliRdel* flagella-less mutants are impaired in swimming motility *in vitro*. (**a**) Schematic representation of the ∼40kb region from the *S. meliloti* 2011 (*Sm* 2011) genome, ranging from *fliF* to *fliR*, that was deleted in the *fliF-fliRdel* deletion mutant to abolish flagella biosynthesis and assembly. The *fliF* deletion (in blue) was generated to abolish flagella assembly. (**b**-**c**) A swimming motility assay was performed by spot-inoculation of the *Smi* 2011-LacZ wild type (WT) strain and derived *fliF* and *fliF-fliRdel* deletion mutants on TY (0.3 - 0.4% agar). Quantification of swimming motility 3-4 dpi (WT n = 40; *fliF* n = 39; *fliF-fliRdel* n = 40) shown in (**b**) along with representative images of bacterial motility (**c**). (**b**) Box plots show the distribution of values (circles) from 3 independent experiments (n=39-40 per sample). First and third quartiles (horizontal box edges), minimum and maximum (whisker tips), median (centerline), mean (solid black triangle), and outliers (crosses) are shown. Letters indicate statistically significant differences between groups (*p* < 2.2e-16, Welch’s ANOVA and Games-Howell test). Scale bars (**c**) = 1 cm. See also Supplemental Table S2.

**Fig. 3.**
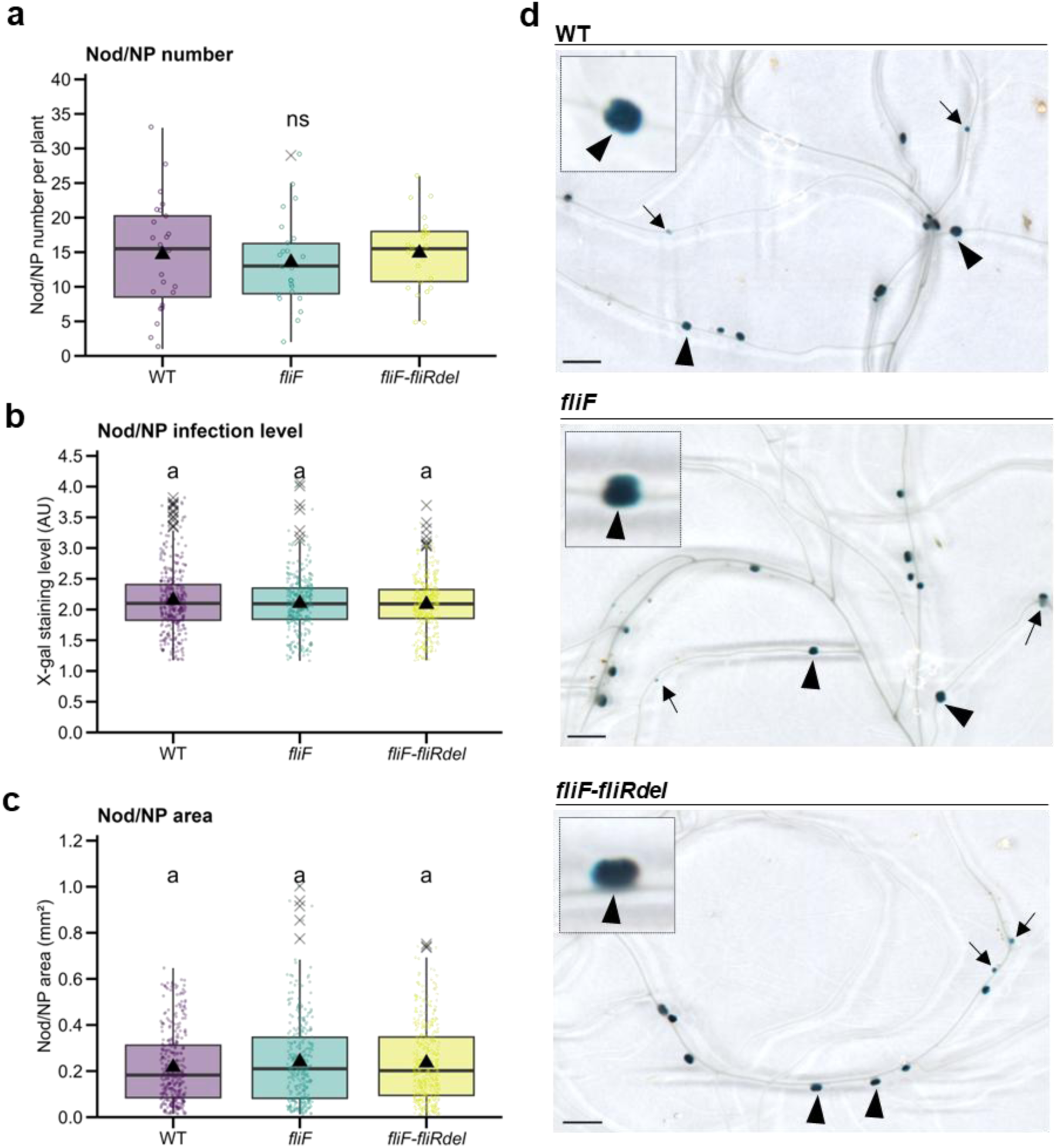
*S. meliloti fliF* and *fliF-fliRdel* flagella motility mutants form nodules which are fully infected in *M. truncatula*. The impact of flagella mutants in early nodule development and infection was quantified in Xgal-stained *M. truncatula* root systems inoculated with *lacZ*-expressing *S. meliloti* WT or *fliF* and *fliF-fliRdel* mutant strains (visualized in blue) at 7 dpi. (**a**-**c**) Number of nodules (Nod) and nodule primordia (NP) per plant (**a**), infection level (X-gal staining intensity) per Nod/NP (**b**), and Nod/NP area (**c**) were quantified at 7 dpi in scanned images of nodulated plants’ root systems (WT n = 24; *fliF* n = 24; *fliF-fliRdel* n = 24 in **a** or individual Nod/NP (WT n = 352; *fliF* n = 325; *fliF-fliRdel* n = 357 in **b**-**c**) from 2 independent experiments. (**d**) Representative images of scanned nodulated root samples. Close-up views of nodules are shown in top left squares. Nodules (arrowheads) and NP (arrows) are indicated. Box plots in **a**-**c** show the distribution of values (circles), first and third quartiles (horizontal box edges), minimum and maximum (whisker tips), median (centerline), mean (solid black triangle), and outliers (crosses). ns (**a**) indicate no statistical difference relative to WT (*p* = 0,766, one-way ANOVA). Classes with the same letter (**b**, **c**) are not significantly different (*p* = 0,6427096 in **b**, *p* = 0,3148374 in **c**, Kruskal-Wallis α = 5%). Scale bars (**d**): = 3 mm. See also Supplemental Fig. S2.

**Fig. 4.**
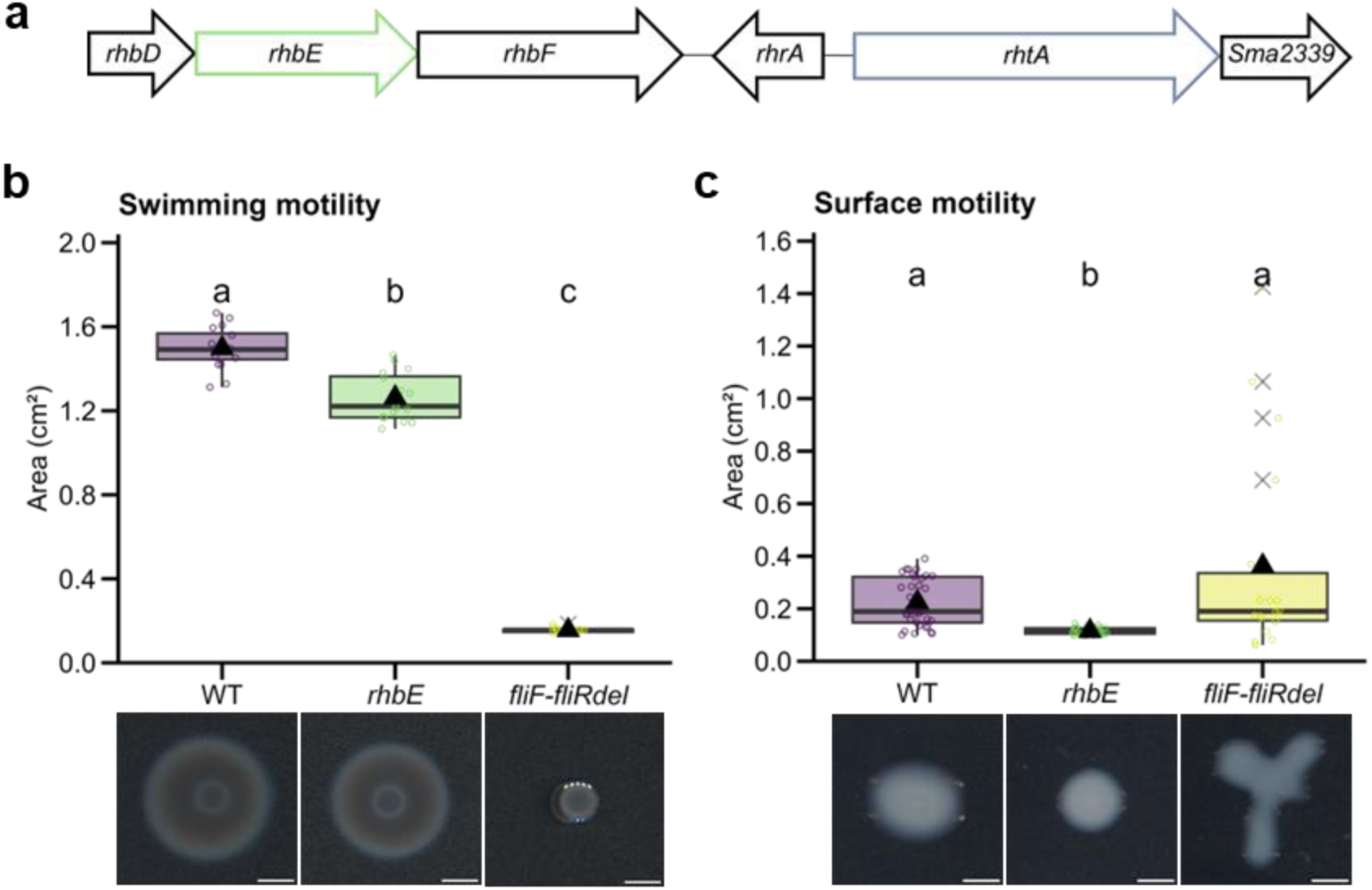
The *S. meliloti* siderophore rhizobactin 1021 biosynthetic gene *rhbE* mutant is impaired in surface motility *in vitro*. (**a**) Schematic representation of the genomic region of *S. meliloti 2011* comprising *rhbE* and *rhtA* genes, which were deleted in the *S. meliloti 2011*-lac Z strain to respectively abolish Rhizobactin 1021 biosynthesis or its utilization. Deleted genes are represented by green and blue boxes. (**b**-**c**) Swimming motility (**b**) and surface motility (**c**) assays were performed by spot-inoculation of *S. meliloti* WT, *rhbE,* and *fliF-fliRdel* strains on plates with TY 0.3% agar (**b**) or MM 0.6% agar (**c**). Box blots show the distribution of values (circles) obtained after ImageJ quantification of bacterial colony growth area 72h after spot inoculation from two (**b**) or three (**c**) technical replicates from 1 (**b**) or 2 (**c**) independent experiments (WT n = 8; *rhbE* n = 8; *fliF-fliRdel* n = 8 in **b**, WT n = 15; *rhbE* n = 12 in **c**). First and third quartiles (horizontal box edges), minimum and maximum (whisker tips), median (centerline), mean (solid black triangle), and outliers (crosses) are shown in box plots. Letters indicate statistically significant differences between groups (*p* < 2.2e-16 in **b**, Welch’s ANOVA and Games-Howell test; *p* = 1.458e-13, Welch two-tailed Student T-test in **c**). Representative images of bacterial growth area on plates are shown below graphs. Scale bars: (**b,c**) = 4 mm. See also Supplemental Table S3.

**Fig. 5.**
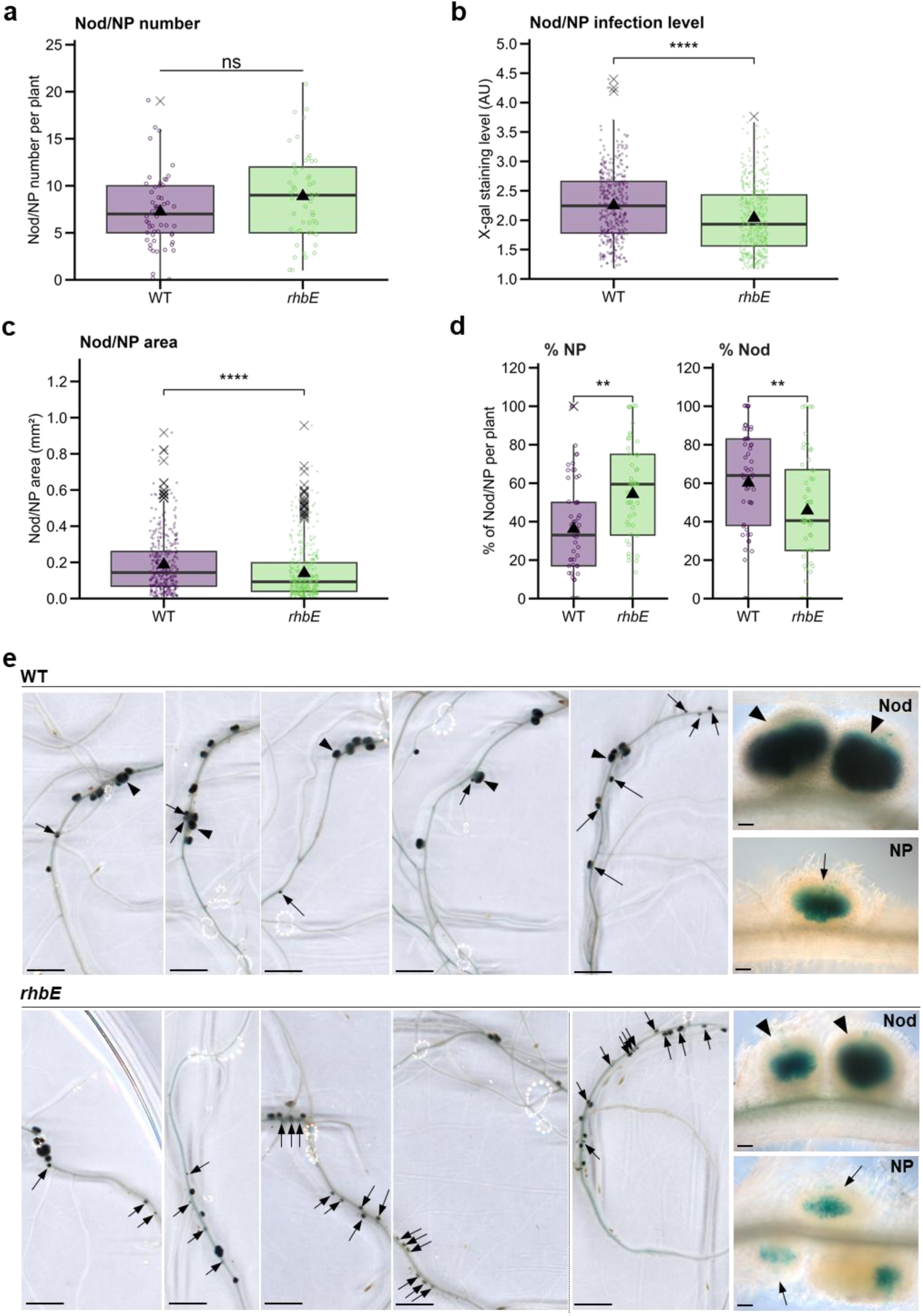
Mutation in the *rhbE* rhizobactin siderophore 1021 biosynthetic gene affects early rhizobia-induced nodule development and colonization. The impact of *rhbE* mutation in early nodule development and infection was quantified in Xgal-stained *M. truncatula* root systems inoculated with *lacZ*-expressing *S. meliloti* WT or *rhbE* mutant strains (visualized in blue) at 7 dpi. (**a**-**c**) Number of nodules (Nod) and nodule primordia (NP) (**a**), Nod/NP infection level (X-gal staining intensity) (**b**) and Nod/NP area (mm2) (**c**) were quantified in individual root systems (WT n = 53; *rhbE* n = 52 in **a**) or individual Nod/NP (WT n = 384; *rhbE* n = 462, in **b**-**c**). (**d**) Percentage of NP (area < 0,100 mm^2^) and Nod (area ≥ 0,100 mm^2^) were quantified per plant (WT n = 53; *rhbE* n = 52). Box plots (**a**-**d**) show the distribution of values (circles) from 3 independent experiments. First and third quartiles (horizontal box edges), minimum and maximum (whisker tips), median (centerline), mean (solid black triangle), and outliers (crosses) are shown. ns (**a**) indicate no statistical difference relative to WT (*p* = 0,05708, two-tailed Student T-test). Asterisks (**b**-**d**) point to statistical differences in *rhbE* relative to WT samples (*p* = 6.389e-08 in **b**, *p* =1.599e-08 in **c**, *p* = 0.001205 for % NP and *p* = 0.00812 for % Nod in **d,** Mann-Whitney test). (**e**) Representative images of *M. truncatula* root systems nodulated with WT or *rhbE* strains. Nod (arrowheads) and NP (arrows) are indicated, and their close-up views are shown in the right corner (**a**-**d**). Scale bars (**e**) = 3 mm; 100 µm (close-up images). See also Supplemental Fig. S2.

### β-galactosidase enzymatic assays

The nodulated roots harvested at 5 or 7 dpi were rinsed in Z buffer (10 mM KCl, 1 mM MgCl_­­_, and 0.1 M phosphate buffer, pH = 7.0) and fixed in a 1.25% (7 dpi) or 2.5% (5 dpi) glutaraldehyde solution for 1h (45 min to 1 h under vacuum) as described previously (Cerri et al., 2012). To reveal the constitutive β-galactosidase activity of the *Sm*2011-lacZ WT and mutant strains, root samples were rinsed twice with Z buffer and stained in Z-buffer containing 2 mM X-gal (5-bromo-4-chloro-3-indolyl-β-D-galactopyranoside, W5376C; Thermo Fisher Scientific, Guilford, CT) at 4°C over weekend or at 28°C overnight. The stained roots were cleared for 1 min with a 12% sodium hypochlorite solution before scanning and microscopic observations.

### Microscopy methods

For analysis of rhizobia cell propagation within ITs (Fig. **1**), *M. truncatula sunn*-2 plants were grown under LUMOX film on phytagel Fahraeus medium and observed from 2 to 5 dpi with *Sm*2011-GFP using a Leica TCS SP2 AOBS laser scanning confocal microscope with a 40x long-distance water immersion objective (HCX APO L U-V-I 40x/0.80 WATER). The argon laser band of 488nm was used to excite GFP and a 561nm diode to excite the cell wall autofluorescence. Specific emission windows used for GFP and autofluorescence signals were 500-530 nm and 620-720 nm, respectively; emitted fluorescence was false colored in green (GFP) and magenta (wall autofluorescence). Merged confocal images and maximum projections of several z-sections were created using the Fiji software for illustrations. The positions of rhizobial cells within ITs were determined by defining Regions of interest (ROIs) and extracting their x,y coordinates using the MicrobeJ plugin of ImageJ.

For experiments in Fig. **6**, *M. truncatula* roots grown under LUMOX film on phytagel Fahraeus medium were observed from 2 to 4 dpi with WT or mutant *Sm*2011-mScarlet using a Leica TCS SP8 AOBS laser scanning confocal microscope with a 40x long-distance water immersion objective (HCX APO L U-V-I 40x/0.80 WATER). A 561nm diode-pumped solid-state (DPSS) laser was used to excite the mScarlet fluorescent proteins, and the emitted fluorescence was detected using a hybrid detector (HyD) in the 585 –640 nm emission window. Leica LAS-X software was used to record confocal images, and the time-series images were then analysed with Fiji software to determine the number of ITs in each RH. Merged confocal images and maximum projections of several z-sections were created using the Fiji software for illustrations.

**Fig. 6.**
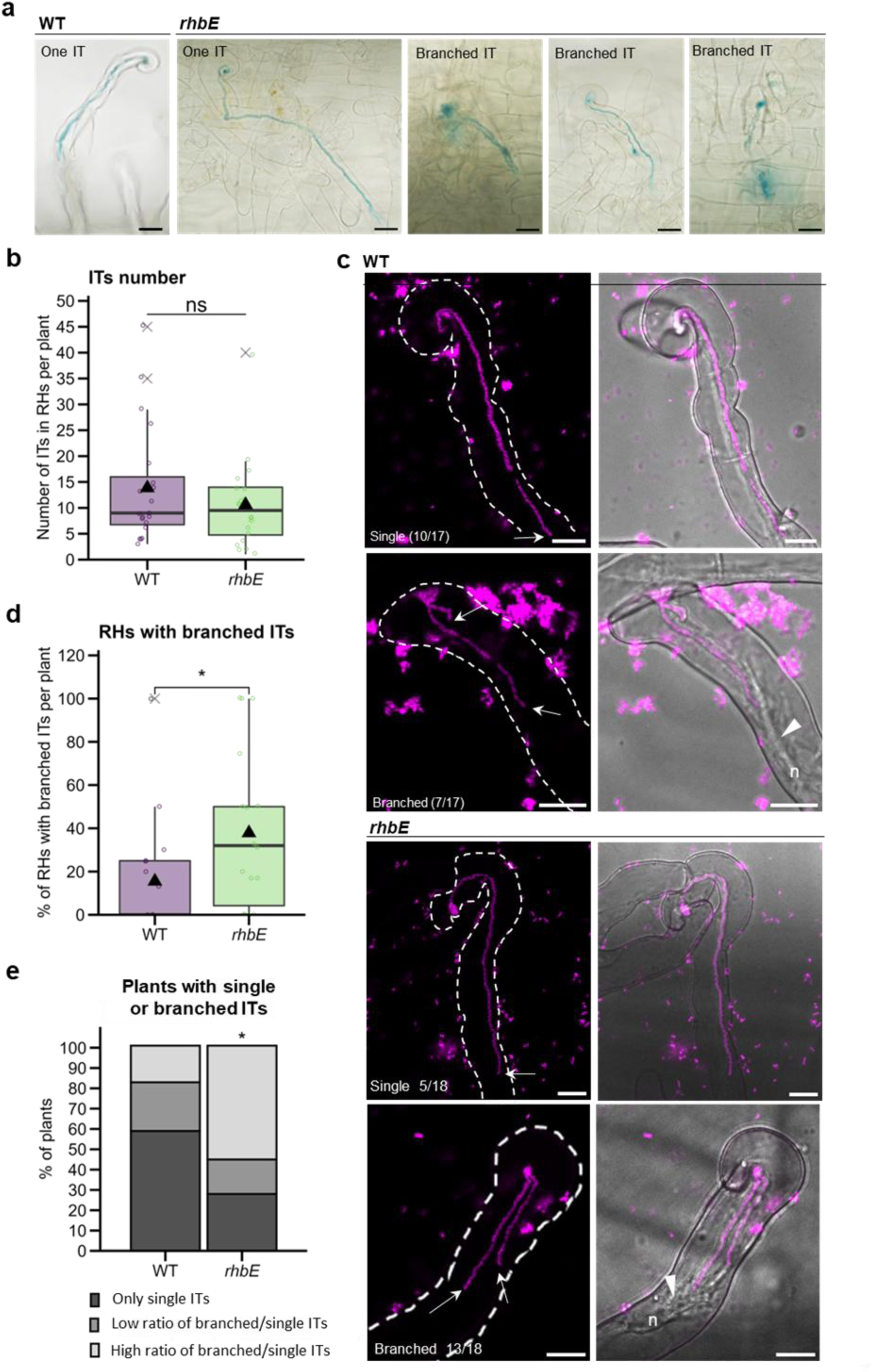
Impact of *rhbE* mutation in IT root hair development in *M. truncatula*. The impact of *rhbE* mutation in early development of ITs in root hairs was analysed in *M. truncatula* Xgal-stained root systems (**a**, **b**) after inoculation with *lacZ*-expressing *S. meliloti* WT or *rhbE* mutant strains (visualized in blue) or *in vivo* (**c**-**e**) in roots inoculated with *S. meliloti* WT or *rhbE* mutant strains expressing a mScarlet fluorescent reporter. (**a**, **b**) Number of ITs in root hairs was quantified by counting root hair lacZ-stained rhizobia infection events (illustrated in **a**) WT and mutant samples on bright field microscopy images taken from slide-scanned rhizobia-inoculated root systems at 5 dpi from 2 independent experiments (WT n = 20; *rhbE* n = 20) (**b**). (**c**-**e**) Confocal microscopy analysis of early IT development using mScarlet *S. meliloti* WT or *rhbE* mutant strains was carried out *in vivo* from 2 to 4 dpi. (**c**) Representative confocal fluorescent and merge images of RHs with single or branched ITs (visualized in magenta). Arrows indicate IT tips, which are linked to the nucleus (n) by a cytoplasmic bridge (white arrowhead). Proportion of RHs with branched ITs per plant (**d**) or of individual plants having only single ITs or having different ratio of branched vs single ITs (**e**): low (≤ 1:4 branched vs single ITs) or high (≥ 1:3 to 1:0 branched vs single ITs). Data in **d**-**e** are from 4 independent experiments (WT n = 17; *rhbE* n = 18). Box plots in **b**, **d**, **e** show the distribution of values (circles) from 2 (**b**) or 3 (**c**) independent experiments. First and third quartiles (horizontal box edges), minimum and maximum (whisker tips), median (centerline), mean (solid black triangle), and outliers (crosses) are shown. ns (**b**) indicate no statistical difference relative to WT (*p* = 0,2525, two-tailed Student T-test). Asterisks (**d, e**) point to statistical differences in mutant relative to WT (*p* = 0,03273 in **d**, Mann-Whitney test; *p* = 0,0354 for 26-100% in **e**, two-tailed Fisher’s exact tests). Scale bars: (**a**) = 20 µm; (**e)** = 10 µm.

For nodulation phenotyping of plants inoculated with *Sm*2011-lacZ WT or mutant strains, β-galactosidase-stained nodulated roots harvested at 7 dpi were scanned (Objectscan 1600, Microtek). The scanned images were used to analyse the nodule numbers and measure the nodule size and infection levels of the β-galactosidase-stained nodulated roots, using the Fiji software. Detection of blue β-galactosidase-stained nodules was performed with the assistance of machine-learning recognition with Fiji software (color threshold method). Each nodule or nodule primordium was described as a separate ROI and counted for each plant. Specific nodule features were measured for each ROI, such as the area and level of infection (based on the blue intensity correlated to the rhizobial β-galactosidase activity). A light microscope (AxioPlan II Imaging; Carl Zeiss, Oberkochen, Germany) was used for illustration of the different categories of Nod/NPs. While β-galactosidase-stained roots harvested at 5 dpi were scanned between sealed slides and coverslips with a 20X objective (NanoZoomer 2.ORS, Hamamatsu Photonics France) at a thickness of 15 histological sections spaced 10µm apart, starting at a depth of 4µm. The ITs were then counted for each root on these scans and photographed for figure illustrations using NDP.view2 software.

For nodulation phenotyping of β-galactosidase-stained nodulated roots in supplementary Fig. **S2**, the nodules were counted using a binocular magnifying glass and divided into 2 categories as follows: infected or uninfected nodules/nodule primordia structures.

### Graphs and statistical analyses

Graphical representation and statistical analyses of the data were performed using R. Data are represented either as box plots or stacked bar charts. The normal distribution of the data was assessed using the Shapiro-Wilk test, and in the case of a non-normal data distribution, a transformation was performed to normalize the data distribution if possible (Log10 or BoxCox). In the case of normal distribution, homogeneity of variance was evaluated using Fisher or Bartlett tests. For normal data distribution, data were analysed with parametric statistical tests (t-test, one-way ANOVA, Tukey test, Welsh’s ANOVA, and Games-Howell test), whereas non-parametric tests (Mann-Whitney, Kruskal-Wallis, and Dunn test) were used for data with a non-normal distribution. Fisher’s tests were used to analyse data in contingency tables. Figure legends indicate the number of replicates and significance p-value levels, as well as the total number of individuals analysed (n). In detail for each figure, the data were analysed as follows: Values in Fig. **2b** follow a normal distribution and the variances were not homogeneous, so a Welch’s ANOVA followed by a Games-Howell test was performed (*F* = 164.17, *p* < 2.2e-16). Values of Fig. **3a** follow a normal distribution and variance homogeneity, so a one-way ANOVA was carried out (*F* = 0.268, *p* = 0.766). In Fig. **3b**, values do not follow a normal distribution and were thus analysed using a Kruskal-Wallis test (*K* = 0.8841246, *p* = 0.6427096). Values in Fig. **3c**, do not follow a normal distribution; hence, a Kruskal-Wallis test was carried out (*K* = 2.311398, *p* = 0.3148374). Values in Fig. **S2a** and b follow a normal distribution and variance homogeneity, so a one-way ANOVA was performed (*F* = 0.113, *p* = 0.894 in a; *F* = 0.225, *p* = 0.799 in b). Values in Fig. **4b** follow a normal distribution and the variances were not homogeneous, so a Welch’s ANOVA followed by a Games-Howell test was performed (*F* = 2048.2, *p* < 2.2e-16). Values in Fig. **4c**, do not follow a normal distribution; hence, a Kruskal-Wallis test was carried out (*K* = 29.35631, *p* = 4.220439e-07). BoxCox-transformed values (λ = 0.5858586), in Fig. **5a**, show normal distribution and variance homogeneity and thus were analysed using a two-tailed Student T-test (*T* = 1.9243, *p* = 0.05708). Values in Fig. **5b**,**c** and d do not follow a normal distribution, so Mann-Whitney tests were applied (*W* = 107840, *p* = 6.389e-08, for Fig. **5b**; *W* = 108700, *p* = 1.599e-08, for Fig. **5c**; *W* = 873, *p* = 0.001205 for % NP and *W* = 1791, *p* = 0.00812 for % Nod, for Fig. **5d**). Values in Fig. **S3**, do not follow a normal distribution; hence, a Kruskal-Wallis test was carried out (*K* = 23.95754, *p* = 6.276046e-06). BoxCox-transformed values (λ = 0.02020202), in Fig. **6b**, show normal distribution and variance homogeneity; hence, a two-tailed Student T-test (*T* = - 1.1619, *p* = 0.2525). In Fig. **6d**, values do not follow a normal distribution and were thus analysed using a Mann-Whitney test (*W* = 90.5, *p* = 0.03273) and confirmed by a permutation test (*p* = 0.0461), which reinforces the reliability of this conclusion despite the presence of ties in the data. Fig. **6e** was created from a contingency table reporting the relative proportion of individual plants having only single ITs or having a different ratio of branched vs single ITs (**e**): low (≤ 1:4 branched vs single ITs) or high (≥ 1:3 to 1:0 branched vs single ITs). Fisher’s exact tests allowed us to compare the relative proportions of these categories between the WT and *rhbE* mutant strain (*p* = 0.0922 for single ITs; *p* = 0.691 for low ratio; *p* = 0.0354 for high ratio).

### Mathematical model of infection thread growth

The cell proliferation and motion within the infection thread was modelled using agent-based modelling mimicking autonomous individual cells with heterogeneous behaviour, in a spatially defined environment, i.e. the Infection Thread, and changing over time. The model is described in detail in the Supplementary data and in brief in the following. Each cell has its own size represented by an ellipse described by a variable length in μm and a fixed width of 0.4 μm, and can divide when it reaches a maximal length at a certain growth rate in h^-1^. The motion of the cell depends on 1) of the migration direction, i.e. the angle in a 2D plane toward which the bacteria move through the infection thread, and 2) the speed in μm/h of the cell motion. To simulate the influence of the environment on the cell behaviour, a 2D grid was created. Each grid cell corresponds to the local micro-environments encountered by bacteria and may have two properties, 1) the resistance of the environment against bacterial motion due for instance to local modification of the plant cell wall properties or variation in viscosity due to the secretion of exopolysaccharides and 2) The resources available in the local environment and that may be used by the bacteria to pay the cost of proliferation and motility. The model proceeds by calculating for each 10 minutes time point the following 4 steps iteratively: step 1 the motion of the cells based on their individual velocity, step 2 then the growth of the cell based on their individual growth rate, step 3) the division of the cell if their size and resource allowed it, step 4) resolving overlap between cells to push overlapping cells forward. The model was used to test various hypotheses by comparing predictions with experimental measurements.

## Results

### IT infection kinetics and modelling suggest slow-motion motility of *S. meliloti* cells in elongating ITs

Previous microscopy studies suggested that elongation of the rhizobia bacterial file inside ITs likely involves slow movement modes (Gage, 2002; Fournier *et al*., 2008). To quantitatively assess the motility speed used by rhizobia inside ITs and to test possible associated motility mechanisms, we built a multi-agent mathematical model mimicking the motion and proliferation of rhizobia bacterial cells. Hence, the contribution of different motility mechanisms like swimming (active) or swarming (passive) can be decoupled from the division-based motility, i.e. sliding (see Table **S5**). To this end, we used a series of 3-5 successive images, acquired at intervals of 40-100 minutes, of individual developing ITs (n = 5). Thanks to the localization of individual bacterial cells over time (see in Fig. **S5**), we could calculate the distances travelled by each bacterium overtime and formulate modelling hypotheses (Fig. **1**). The model was parametrised with the shape of the IT and position of the bacterial cells using bright field and confocal microscopy images and fit to experimental data to infer contribution of various types of motilities. Analysis of the experimental data, thanks to the model, revealed that the cells at the forefront of the IT are more mobile compared to those backwards, in accordance with Gage (2004). An average speed of 3.8 ± 1.7 µm per hour was found for the forefront 6 cells in each experiment, and the cells dividing harbour a doubling time with a minimal value of 3.6 hours. Thus, a maximal motion of 0.48 µm per hour can be due to cell division, i.e. sliding motility. This level represents around 15% of the speed of the most mobile cells. Then, the sliding-independent motility may represent around on average, 3.3 µm per hour (85%). The order of magnitude of the speed of the cells is around 4 µm per hour, and between 2 and 6 µm per hour.

We compared the sliding-independent motility contribution (∼4 µm per hour) with previously published speeds of bacterial motility (Table **S5**). In liquid or semi-solid media, flagella-dependent motility (*i.e.* swimming and swarming) propel bacterial cells at very high speeds (5-10 mm per hour) (Table **S5**) up to 3 orders of magnitude higher than the speed observed in ITs. It is possible that flagella-dependent motility is active in ITs, but the resistance within the IT lumen due to high viscosity or friction may be responsible for the much lower speed of motion observed. However, since the motility in a liquid environment is estimated to consume up to 2-3,4% of the cell energy (Schavemaker & Lynch, 2022), it is highly unlikely that swimming inside the IT would be possible. Considering this discrepancy in speed, we hypothesized that flagella-independent motility, including sliding (Nan & Zusman, 2016), might be involved in the progression of rhizobia within early ITs.

### *S. meliloti fliF* and *fliR-fliRdel* flagella mutants are impaired in swimming motility *in vitro*

To investigate the role of flagella-dependent motility during host infection (i.e. swimming and swarming), the 39,838 bp chromosomal flagellar regulon region, extending from *fliF* (encoding the flagellar MS-ring rotor protein) to *fliR* (encoding a probable flagellar biosynthesis protein) (Sourjik *et al*., 2000) was deleted in the genome of *S. meliloti* 2011 (*Sm*2011), resulting in the *fliF-fliRdel* markerless mutant strain (Fig. **2a**). A second deletion, focused on *fliF* (in blue in Fig **2a**), was also created to specifically abolish flagella assembly. Genome sequence comparison of the reference *Sm*2011 wild type (WT) and the *fliF-fliRdel* strain confirmed the deletion of the flagellar regulon region (Fig. **2a**) (Table **S6**).

To later facilitate histochemical visualization of the strains during *in planta* infection, a plasmid constitutively expressing a β-galactosidase (*lacZ*) reporter (pXLGD4, Leong et al.,1985) was introduced in these strains, which are hereafter referred to as *Sm*2011*-lacZ* WT, *fliF* or *fliF-fliRdel.* To address their swimming abilities, swimming motility tests were performed on 0.3-0.4% agar TY plates by measuring the colony migration growth zone at 3-4 dpi. The WT strain spread through the medium, covering an area approximately four times larger than that of *fliF* and *fliF-fliRdel* mutants (Fig. **2b**-**c**). These results confirm the swimming ability of the WT strain over time and show that swimming motility is abolished in the *fliF* and *fliF-fliRdel* strains *in vitro*.

In summary, these data demonstrate that both flagella mutants exhibit equivalent impairments in flagella-dependent swimming motility *in vitro,* regardless of whether the deletion affects a single gene responsible for flagellum assembly (*fliF*) or the entire flagellar regulon (*fliF-fliRdel*).

### *S. meliloti fliF* and *fliF-fliRdel* flagella motility mutants do not exhibit early nodule development and colonization phenotypes *in planta*

While the motility of rhizobia in free-living conditions has been extensively studied (Gotz & Schmitt, 1987; Nogales *et al*., 2012; Wadhwa & Berg, 2022), the specific form of motility they employ to navigate inside the confined IT environment remains unknown. The motility model outlined in Fig. **1** suggested minimal contribution of flagella-dependent motility inside ITs. To investigate this, we compared the ability of lacZ-tagged *fliF* and *fliF-fliRdel* flagella-less mutants to induce and colonize nodules on *M. truncatula* roots relative to the *Sm*2011*-lacZ* WT control. By improving a recently developed method (Guillory *et al*., 2024) based on Image J quantification of X-gal-stained root systems infected with *lacZ*-expressing *S. meliloti* strains, we were able to monitor the number and size of nodules (Nod) and nodule primordia (NP) as well as their infection level in root systems of individual plants. A similar number of nodules (Nod) and nodule primordia (NP) were formed in plants inoculated with flagella-less mutants compared to the WT control (Fig. **3a**). Moreover, nodules and NP formed with flagella-less mutants were well infected and displayed no obvious signs of developmental defects compared to the WT control (Fig**. 3b**-**d**). Similar conclusions were reached when plants were grown under another growth condition and quantified via non-automated, user-based microscopy counting (Fig. **S2**).

In conclusion, flagella-less mutants do not exhibit significant differences in nodule formation or infection levels in *M. truncatula* compared to the control strain when flood-inoculated, regardless of growth conditions or quantification methods. These results are consistent with the model (Fig. **1**) and previous studies in other legume species (Ames & Bergman, 1981; Mellor *et al*., 1987).

Mutation in the *rhbE* rhizobactin siderophore 1021 biosynthetic gene affects *S. meliloti* surface motility *in vitro* and *in planta* root nodule colonization.

The model (Fig. **1**) and experimental data obtained with flagella-less mutants (Fig. **2** and **S2**); both support that *S. meliloti* uses flagella-independent motility modes to colonize the roots of its host plant. *S. meliloti* can engage in flagella-independent sliding surface motility, which relies on the secretion of surfactant-like compounds, such as EPS II and Rhb1021, to reduce surface friction (Nogales *et al*., 2012). As the *S. meliloti* strain 2011 used in this study is impaired in EPS II production (Nogales *et al*., 2012), we focused here on evaluating the role of Rhb1021 in the symbiotic interaction. Mutations in *rhb* genes were previously shown to block the synthesis of Rhb1021 and to render *S. meliloti* 2011 nonmotile on semisolid MM (Lynch *et al*., 2001; Nogales *et al*., 2010, 2012). Here, we created a markerless deletion of *rhbE* (Fig. **4a**) using homologous recombination, which was subsequently confirmed by genomic sequencing (Table **S7**). Unlike the flagella-less *fliF-fliRdel* mutant, the *rhbE* mutant could still swim in TY agar 0.3 % plates comparable to the WT control at 72 hpi (Fig. **4b**). Conversely, *rhbE* showed significant impairment of surface translocation on semisolid MM plates compared to WT and *fliF-fliRdel* strains (Fig. **4c**). Together, these results confirm the involvement of Rhb1021 in promoting passive surface motility of *S. meliloti*.

To investigate how impairment in Rhb1021 production could impact root nodulation, we examined the ability of the *rhbE* mutant strain (also expressing *LacZ*) to colonize and develop nodules in *M. truncatula* relative to the WT control, using the same ImageJ quantification method as used with the flagella mutants (Fig. **3**). At 7 dpi, the total number of nodules (Nod) and nodule primordia (NP) formed per root system with the *rhbE* strain was comparable to those formed with the WT strain (Fig. **5a**). However, infection levels and overall sizes of *rhbE* Nod and NPs (Fig. **5b**-**c**) were consistently reduced compared to those in WT plants. A detailed analysis of Nod and NP distribution per individual plants revealed the overrepresentation of small and underinfected NPs (arrows) in *rhbE* and the opposite overrepresentation of larger well well-infected nodules (arrowheads) in WT-inoculated plants (Fig. **5d**-**e**). Together, these results suggest that Rhb1021 biosynthesis by rhizobia contributes to root nodule colonization.

As secreted siderophores can also function as high-affinity iron chelators to acquire iron when it is scarce in the environment (Timofeeva *et al*., 2022), it remained to be established if the *in-planta* phenotype of the *rhbE* strain was due to a defect in rhizobactin-mediated sliding motility or rather in iron uptake. To distinguish between these possibilities, we generated a markerless deletion of *rhtA* encoding the Rhb1021 outer membrane receptor (Fig. **4a**), which is expected to affect Rhb1021 utilization to scavenge iron but not the ability to synthesize it (Lynch *et al*., 2001). This markerless deletion was subsequently confirmed by genomic sequencing (Table **S8**). As in previous analyses (Fig. **5c**), Nod and NPs in roots inoculated with *rhbE* were significantly less infected (Fig. **S3**). In contrast to the *rhbE* mutant, nodules and NPs formed in *rhtA*-strain inoculated roots showed similar infection levels to those formed in WT-inoculated roots (Fig. **S3**). Collectively, these findings imply that early defects in nodule development and infection by the *rhbE* Rhb1021 biosynthetic mutant are likely attributable to a defect in sliding motility.

### The *rhbE* mutation favours the development of branched ITs in-root hairs

To further dissect the observed *rhbE* mutant infection phenotype, we closely inspected early root hair infection events driven by *rhbE* compared to the WT strain. Using rhizobia strains expressing the *lacZ* reporter, ITs formed in *M. truncatula* root hairs 5 dpi were visualized and quantified (Fig. **6a**-**b**). The number of ITs formed per root hair was not significantly different between the two strains (Fig. **6b**). Nevertheless, RHs with branched ITs (two or three branches) per root hair were frequently observed in root systems inoculated with the *rhbE* strain (Fig. **6a**). To better quantify these early infection events, we used WT and *rhbE* strains constitutively expressing an mScarlet fluorescent reporter for visualizing fluorescent-labelled ITs in root hairs 2 to 4 dpi using *in vivo* confocal microscopy (Fig. **6c**-**e**). Growing ITs, recognized by the presence of a cytoplasmic bridge (arrowhead) connecting the nucleus (n) to the IT tip (arrow), were imaged in both *rhbE* and WT-inoculated plants (Fig. **6c**) for subsequent ImageJ-based quantification. Roots inoculated with the *rhbE* strain showed preferential formation of branched ITs per RH compared to the WT control (Fig. **6d**). Furthermore, a higher proportion of branched ITs was found in individual *rhbE* plants than in the WT control (Fig. **6c**, **d**). Indeed, most WT plants had only single ITs (60 %, 10/17), whereas this category represented only 30 % of *rhbE* plants (5/18). Similar proportions of plants (24% in WT, 4/17 and 17 % in *rhbE*, 3/18) exhibited low ratios of branched ITs (≤ 1:4 branched vs single ITs). In contrast, plants with a high proportion of branched ITs (1:3 to 1:0 branched vs single ITs) were overrepresented for the *rhbE* strain (10/18) compared to WT (3/17). Together, these results show that the *rhbE* mutation partially impacts the development of ITs in root hairs, which may explain the observed reduced host colonization. Mutation of *rhbE* strikingly promotes IT branching in RHs, raising the question of the need of Rhb1021 for guiding tip directional growth of ITs.

## Discussion

In this study, we integrated mathematical modelling, live cell imaging, and *in planta* phenotypic analyses of bacterial mutants with reduced motility to elucidate privileged motility modes used by rhizobia during early stages of colonization of their legume host. Focusing on the *S. meliloti*-*M. truncatula* symbiotic model system, our findings indicate that flagella-less *S. meliloti fliF* and *fliF-fliRdel* mutants, which are abolished in swimming motility *in vitro*, are not impaired in nodule initiation, development, or infection (Fig. **2**-**3**). In contrast, an *S. meliloti rhbE* mutant, which is blocked in Rhb1021 biosynthesis and affected in surface motility (Nogales *et al*., 2010, 2012 and this work, Fig. **4**-**6**), can initiate nodule primordia and nodule formation, but these are affected in their development and infection level. These findings are consistent with the estimated model (Fig. **1**) that rhizobia preferentially rely on passive flagella-independent paths to colonize their host and provide new evidence that siderophore Rhb1021 biosynthesis by rhizobia plays a role in legume host infection.

Here, we provided a detailed phenotypic characterization and quantification of early infection and nodule development phenotypes of flagella-less mutants *fliF* and *fliF-fliRdel*, not done before in rhizobia-legume symbiosis. The lack of an obvious symbiotic phenotype of flagella-less mutants *fliF* and *fliF-fliRdel* in *M. truncatula* when bacteria are flood inoculated (Fig. **3**) is in line with previous studies in other rhizobia-legume symbiotic interactions showing that flagella are not essential for late nodule development (Ames & Bergman, 1981; Mellor *et al*., 1987; Salas *et al*., 2017; Navarro-Gómez *et al*., 2024). While motile strains are more competitive for nodule occupancy compared to non-motile strains (reviewed in Aroney *et al.,* 2021), this advantage likely comes at earlier stages, such as movement toward the root (Mellor *et al*., 1987; Bernabéu-Roda *et al*., 2015; Navarro-Gómez *et al*., 2024), attachment (Fujishige *et al*., 2006; Zheng *et al*., 2015), and spreading on the root surface (Caetano-Anollés *et al*., 1988). Nevertheless, a transposon insertion sequencing genetic study showed that flagella-less mutants are impacted in their survival and growth at later stages in nodules (Wheatley *et al*., 2020). Though our results do not support a prominent role of flagella during early host colonization, high-resolution *in vivo* biotracking tools (Ozer *et al*., 2021) could provide complementary relevant information on when flagella are actually lost during the symbiotic colonization of the host.

Our experimental data and model (Fig. **1** and **3**) suggest that the collective movement of rhizobia through the IT compartment can occur independently of flagella. Our model favours slow (2 to 6 µm/h) over rapid (5 mm/h) (Table **S5**) flagella-dependent movement within early root hair IT compartments. This is consistent with the observation that at high cell population densities, as inside ITs, genes associated with the flagellar regulon are repressed (Gurich & González, 2009). Building on previous observations that the synthesis of the siderophore Rhb1021 contributes to surface motility (Nogales *et al*., 2010, 2012; Bernabéu-Roda *et al*., 2015), compatible with the slow movement predicted by the model (Fig. **1**), we conducted a genetic study of the role of Rhb1021 biosynthesis and uptake in the *S. meliloti*-*Medicago* symbiosis. Phenotypic characterisation of the Rhb1021 biosynthetic *S. meliloti rhbE* mutant provided novel evidence for a role of Rhb1021 in proper development and colonization of NP and nodules in *Medicago*. As infection and nodule development are interconnected processes (Xiao *et al*., 2014), we believe that the nodule size developmental defects observed in *rhbE* inoculated plants might be a consequence of the impaired NP and nodule colonization (Fig. **5**). Defective nodule phenotypes of the *rhbE* strain could possibly impact later stages of nodule differentiation and functioning, which could explain the reduced nitrogen fixation efficiency previously described for another Rhb1021 biosynthesis mutant (Gill *et al*., 1991). As no obvious nodulation phenotypes were observed with the *rhtA* mutant (Fig. **S3**), which can synthesize Rhb1021 but cannot use it to acquire iron, we can confidently conclude that the symbiotic phenotypes observed with *rhbE* are not due to an iron-scavenging issue but rather due to the absence of Rhb1021 per se, which likely permits optimal motility during host infection. These results are consistent with a previous study showing that a *Rhizobium leguminosarum* mutant impaired in siderophore uptake but not biosynthesis was unaffected in its ability to induce N_­­_-fixing nodules in pea (Stevens *et al*., 1999).

In line with previous studies (Nogales *et al*., 2010, 2012), the Rhb1021 biosynthesis *rhbE* mutant produced in this study showed impaired surface motility in semi-solid MM medium (Fig. **4**). As for other surfactants (Burch *et al*., 2012), it has been proposed that Rhb1021 production could be coordinated with flagellar assembly and thus be affected in flagella-less mutants (Bernabéu-Roda *et al*., 2015). Considering the distinct phenotypes of flagella and *rhbE* mutants *in vitro* (Fig. **2** and **4**) and *in planta* (Fig. **3** and **5**), this is likely not the case in this strain. Thus, the observed *in planta* phenotype of *rhbE* is likely due to impairment of a flagella-independent motility process.

The impact of impaired surface motility of the *rhbE* mutant on host infection likely did not arise from a defect in reaching root hairs, since the roots were flood-inoculated similarly to the flagella mutants. It is possible that the *rhbE* mutant is impaired in attachment or spreading on the root surface, as seen in a previous study (Amaya-Gómez *et al*., 2015). However, it has been shown that spreading on the root surface primarily occurs passively through root elongation, not bacterial motility (Caetano-Anollés *et al*., 1988). Furthermore, no difference was observed in the number of ITs formed in root hairs by the *rhbE* mutant (Fig. **6b**), suggesting that the symbiotic defects of *rhbE* are only manifested after the initiation of infection in our experimental system. The *rhbE* mutant strain strikingly promotes IT branching in root hairs (Fig. **6**), raising the question of the need of Rhb1021 for normal IT development. It is possible that Rhb1021 secretion to the infection chamber somehow regulates the oriented tip growth of the IT. Recent cytochemical studies suggested ROS enrichment of the IT to possibly regulate cell wall stiffness during IT growth (Tsyganova *et al*., 2024). As siderophores have been associated with ROS sequestering or production (reviewed in Arnold, 2024), there may be a connection between Rhb1021 and ROS-mediated cell wall modifications for IT growth. It would be interesting to investigate how an *rhbE* mutation would affect ROS homeostasis or other cellular mechanisms of IT growth (Jamet *et al*., 2003, 2007; Puppo *et al*., 2013; de Carvalho-Niebel *et al*., 2024).

Overall, our data suggest that *S. meliloti* uses flagella-independent surface translocation through the secretion of the surfactant Rhb1021 to fine-tune directed IT tip growth for optimal colonization of developing nodules of *M. truncatula.* Though *rhbE* mutation impacts proper nodule colonization and root hair IT development, ITs can still form and progress towards developing nodules. Thus, it is likely that rhizobia motility inside ITs depends on Rhb1021 and other mechanisms. The galactoglucan exopolysaccharide (EPS II) is used by *S. meliloti* to promote surface motility (Nogales *et al*., 2012). However, *S. meliloti* 2011, the strain used in this work, has a mutation in the *expR* gene, which encodes a key activator of EPS II biosynthesis (Pellock *et al*., 2002). *S. meliloti* can also synthesize succinoglycan (EPS I) (Cheng & Walker, 1998) with major roles in early rhizobia infection signalling (Acosta-jurado *et al*., 2021). Like other surfactants, EPS I biosynthesis can also promote surface motility in *S. meliloti*, though this has only been shown in overexpression conditions (Nogales *et al*., 2012). Future genetic or live microscopy studies may help elucidate the expression of EPS I biosynthesis genes during host infection and the potential involvement of this compound in rhizobia motility inside the IT.

Finally, slow motility requires surfactants but also may involve force generated from cell proliferation. Our model and live imaging in early IT formation suggest that the force generated by proliferation may represent an order of magnitude of around 15% of all mechanisms responsible for the bacterial motion. This observation differs from Gage modelling and observations made with mature ITs, where sliding motility was suspected to be the main mechanism of motility. An investigation of the spatiotemporal dynamics of bacterial cell proliferation during host infection could therefore further expand our understanding of how rhizobia move through the IT.

## Acknowledgements

This work was supported by the French National Research Agency ANR (grants ANR-19-CE20-0026-01 and ANR-24-CE20-6206) and the German Research Foundation DFG (grant BE 2121/9-1). AD was funded by a PhD grant from the French Ministry of National Education and a TULIP-GS Side Project grant (ANR-10-LABX-4). AB^1^ by DFG (grant BE 2121/9-1) and a Binational Doctoral Candidate Motility Grant from Philipps Universität Marburg. AG was funded by an ANR postdoctoral grant (ANR-19-CE20-0026-01) and LM by ENSA3. We would like to thank the Toulouse TRI-FRAIB Imaging platform and especially Aurélie Le Ru, who improved the ImageJ macro for nodule counting; Heiko Wendt, for providing the Sm2011 *fliF* deletion mutant; and Ludovic Legrand, from the LIPME Bioinformatics platform for his help with bacterial genome assembly. We also would like to thank Laurent Sauviac and Fiona Ullmann for their advices on bacterial genomic DNA extraction and Delphine Capela for her helpful advice on surface motility tests.

## Author contributions

AB^2^ and FdCN conceived and supervised the study. JF generated confocal images for the modelling studies that were performed by FB and RP. AB^1^ generated and validated the strains with the help of EK. AB^1^, AG, and LF performed the in vitro motility tests. AD did the mutant phenotypic studies in *Medicago* with the help of AG and LF. LM set up the slide scanning system for in planta infection quantification. JS mapped and analysed mutant genomic data. AD, AB^1^, AG, JF, LF, JS, FdCN, AB^2^ analysed and interpreted the experimental data. AD, AB^1,^ and FdCN wrote the paper with inputs from AB^2^, RP, JF, and AG.

## Data availability

The authors declare that all data supporting the results of this study are available in the article and in Supporting Information Files. Materials generated in this study, including the ImageJ macro used for quantification of nodule size and X-gal staining, are available from the corresponding author upon request. Genome sequence data are available under accession numbers xxx (as placeholder xxx).

