## Supplemental Information for "Rhizobial motility preference in root colonization of *Medicago truncatula*"

**Supporting Information, manuscript Delers & Bennion et al.**

### Tables:

### **Table S1.** Bacterial strains and plasmids.

| **Strains or plasmids** | **Properties** | **Reference** |
|---|---|---|
| ***E. coli strains*** | | |
| DH5α | *E. coli, recA1*, Δ*lacU169*, Δ80d*lacZ*, ΔM15 | Bethesda research Laboratories |
| XL1-Blue | *E. coli* XL-1 blue MRF', *recA1 lac thi* [F' traD36 *proAB lacI*$^q$ *Z*ΔM15] Tc$^r$ | Bullock *et al.,* 1987 |
| S17-I | *E. coli* 294, *thi*, RP4-2-Tc::Mu-Km::Tn*7* integrated into the chromosome | Simon, 1984 |
| DH5α pAB14 | DH5α with pAB14 plasmid Spec$^r$ | This work |
| S17-I pXLGD4 | S17-I with pXLGD4 plasmid, Tc$^r$ | This work |
| S17-I pAB42 | S17-I with pAB42 plasmid, Tc$^r$ | This work |
| S17-I pAB43 | S17-I with pAB43 plasmid, Tc$^r$ | This work |
| S17-I pAB44 | S17-I with pAB44 plasmid, Tc$^r$ | This work |
| ***S. meliloti strains*** | | |
| *Sm*2011 WT | *S. meliloti* 2011 wild type, *expR*$^-$, Strep$^r$ | Casse *et al.*, 1979 |
| *Sm*2011-lacZ | *Sm*2011 WT with pXLBD4 plasmid, Strep$^r$ Tc$^r$ | Cerri *et al.*, 2012 |
| *Sm*2011-GFP | *Sm*2011 WT with pHc60 plasmid, Tc$^r$ | Fournier *et al.*, 2015 |
| *Sm*2011-mScarlet | *Sm*2011 WT with pAB14 plasmid, Strep$^r$ Spec$^r$ | This work |
| Δ*fliF* | *Sm*2011 with a markerless deletion of *fliF* gene (*SMc03014*) (chr<711589 – 713262>), Strep$^r$ | H. Wendt, SYNMIKRO |
| Δ*fliF*-lacZ | Δ*fliF* with pXLGD4 plasmid Strep$^r$ Tc$^r$ | This work |
| *fliF-fliR*del | *Sm*2011 with markerless deletion of ~40 kb from *fliF* (*SMc03014*) to *fliR* (SMc03055) (chr<711589 – 751436>), Strep$^r$ | This work |
| *fliF-fliR*del-lacZ | *fliF-fliR*del with pXLGD4 plasmid Strep$^r$ Tc$^r$ | This work |
| Δ*rhbE* | *Sm*2011 with a markerless deletion of *rhbE* gene (MpA<1311399 – 1312769>), Strep$^r$ | This work |
| Δ*rhbE*-lacZ | Δ*rhbE* with pXLGD4 plasmid Strep$^r$ Tc$^r$ | This work |
| Δ*rhbE*-mScarlet | Δ*rhbE* with pAB14 plasmid Strep$^r$ Spec$^r$ | This work |
| Δ*rhtA* | *Sm*2011 with a markerless deletion of *rhtA* gene, (MpA<1315745 – 1317985>) Strep$^r$ | This work |
| Δ*rhtA*-lacZ | *rhtA* with pXLGD4 plasmid Strep$^r$ Tc$^r$ | This work |
| ***Plasmids*** | | |
| pABC-Psyn | Mobilizable repABC-based mini-replicon, single-copy in *S. meliloti*, Spec r | Schäper *et al.*, 2019 |
| pK18mobsacB | Suicide vector, pUC18 derivate, *lacZ*α, *mob* site, *sacB,* Km$^r$ | Schäfer *et al.*, 1994 |
| pXLGD4 | pRK290 derivative, *hemA*(*SMc03104*)::*lacZ*, Tc$^r$ | Leong *et al.*, 1985 |
| pAB14 | pABC-psyn-p*SMc06412::mScarlet-I*, Spec$^r$ | This work |
| pAB42 | pK18mobsacB-*fliF* 5'UTR$_{-1565--1}$::*fliR* 3'UTR$_{1-1565}$, Km$^r$ | This work |
| pAB43 | pK18mobsacB-p*rbhE*$_{-709--1}$::*rhbE*::*rhbE* 3'UTR$_{1-714}$, Km$^r$ | This work |
| pAB44 | pK18mobsacB-p*rhtA*$_{-700--1}$::*rhbA*::*rhbA* 3'UTR$_{1-698}$ | This work |

**Table S2.** Synthetic DNA used in this work.

| ID | Name | Sequence |
|---|---|---|
| 1 | mScarlet coding sequence<br><br>Original sequence (Bindels *et al.* 2016) codon optimized for *S. meliloti* and synthesized by Twist Bioscience | ATGGTCAGCAAGGGCGAAGCCGTCATCAAAGAATTCATGCGCTTCAAGGTCCAC<br>ATGGAAGGCTCGATGAACGGCCACGAGTTCGAGATCGAAGGCGAAGGCGAAG<br>GCCGGCCGTATGAGGGCACCCAGACCGCCAAGCTCAAGGTCACCAAAGGCGGC<br>CCGCTGCCGTTCTCGTGGGACATCCTTTCGCCGCAGTTCATGTATGGCTCGCGCG<br>CATTCATCAAGCATCCGGCAGACATCCCGGACTATTATAAGCAGTCGTTCCCGGA<br>AGGCTTCAAGTGGGAGCGCGTCATGAACTTCGAGGATGGCGGCGCAGTCACCG<br>TCACGCAGGACACCTCGCTTGAGGACGGCACCCTCATCTATAAGGTCAAGCTTC<br>GCGGCACGAACTTCCCGCCGGATGGCCCGGTCATGCAGAAAAAGACCATGGGC<br>TGGGAAGCCTCGACCGAGCGGCTCTATCCGGAAGATGGCGTCCTGAAGGGCGA<br>CATCAAGATGGCCCTCCGCCTCAAAGATGGCGGCCGCTACCTCGCCGACTTCAA<br>GACCACCTATAAGGCCAAGAAACCGGTCCAGATGCCGGGCGCATATAACGTCGA<br>CCGCAAGCTCGACATCACCTCGCACAACGAGGACTATACCGTCGTCGAGCAGTAT<br>GAGCGCTCGGAAGGCCGCCATTCGACCGGCGGCATGGACGAGCTGTATAAGTG<br>A |
| 2 | PncA-f | ttaagcggccgcAAATGCAAAAAAAGCAGATGACAGCGCCCAAGGTTTGAGCTAAT<br>TTCGGCGCAAAGTGTCGTTGTGGAAAAGATTCCGATAGGAGAAAGAAATGgcta<br>gcatat |
| 3 | PncA-r | atatgctagcCATTTCTTTCTCCTATCGGAATCTTTTCCACAACGACACTTTGCGCCG<br>AAATTAGCTCAAACCTTGGGCGCTGTCATCTGCTTTTTTTGCATTTgcggccgcttaa |
| 4 | mScarlet-MCS-f | atagcggccgcaagcttagatctatagctagcATGGTCAGCAAGGGCGAGGC |
| 5 | mScarlettSm-Kpn-r | ataGGTACCTCACTTATACAGCTCGTCCATG |
| 6 | rhbE-US-f | atagaattcaTGCAAGGCGAATCTCAAG |
| 7 | rhbE-US-r | ataggtaccCATAATCCGAATTGTTGAATG |
| 8 | rhbE-DS-f | ataggtaccCTCATGCATCATGATCCGC |
| 9 | rhbE-DS-r | atatctagaTGGATGCACCGGCATCATG |
| 10 | rhtA-US-f | atatctagacggCATGTTCCTCGCTCTGA |
| 11 | rhtA-US-r | ataggatcccatAACGCCCCCTGCGCAC |
| 12 | rhtA-DS-f | ataggatccGACCGGGGTGATCGCGGAC |
| 13 | rhtA-DS-r | tctaagcttGGATCGGCGAACATTCGCTC |
| 14 | flaXX-HR1-XbaI-f | atatctagaTATGCCCTGACAGGCGTCGAG |
| 15 | flaXX-HR1-NotI-r | atatgcggccgcTCTGTTTCCGCACCCAGTAT |
| 16 | flaXX-HR2-NotI-f | atatagcggccgcGCTCGATGAAGGCGCGGT |
| 17 | flaXX-HR2-HindIII-r | gcagaagcttCAGAAATCGCTGTAGCGGAA |

For primers, annealing regions are indicated with uppercase letters and appendix sequences added for cloning purposes are indicated with lowercase letters.

### **Table S3.** Plasmid construction.

| Plasmid | Construction process |
|---|---|
| pAB14 | A 50 bp region of the *SMc06412* promoter and a 44 bp *mucR* leader and Shine Dalgarno sequence and its reverse complement (oligonucleotides 2 and 3, Table S4) were hybridized and digested with NotI and NheI. The mScarlet coding sequence (oligonucleotide 1, Table S4) was PCR-amplified (oligonucleotides 4 and 5, Table S4) and digested with NheI and KpnI. Inserts were ligated into the pABC-Psyn vector opened with NotI and KpnI and dephosphorylated with FastAP to prevent recyclization. |
| pAB42 | A ~1,5 kb region upstream of *fliF* was PCR-amplified from Rm2011 genomic DNA (oligonucleotides 14 and 15, Table S4) and digested with XbaI and NotI. Likewise, a ~1,5 kb region downstream of *fliR* was amplified (oligonucleotides 16 and 17, Table S4) and digested with NotI and HindIII. These inserts were ligated into the pK18mobsacB vector opened with XbaI and HindIII and dephosphorylated with FastAP to prevent recyclization. |
| pAB43 | A ~700 bp region upstream of *rhbE* was PCR-amplified from Rm2011 genomic DNA upstream (oligonucleotides 6 and 7) digested with EcoRI and Acc65I. Likewise, a ~700 bp region downstream of *rhbE* was amplified (oligonucleotides 8 and 9) and digested with Acc65I and XbaI. These inserts were ligated into the pK18mobsacB vector opened with EcoRI and XbaI and dephosphorylated with FastAP to prevent recyclization. |
| pAB44 | A ~700 bp region upstream of *rhtA* was PCR-amplified from Rm2011 genomic DNA (oligonucleotides 10 and 11) and digested with XbaI and BamHI. Likewise, a ~700 bp region downstream of *rhtA* was amplified (oligonucleotides 12 and 13) and digested with BamHI and HindIII. These inserts were ligated into the pK18mobsacB vector opened with XbaI and HindIII and dephosphorylated with FastAP to prevent recyclization. |

### **Table S4.** Genome sequencing of parental *S. meliloti* WT and mutant strains.

| Strain | Sequencing method | Total bp sequenced | Total reads | Raw coverage | Assembled coverage |
|---|---|---|---|---|---|
| 2011 WT (parental strain) | ONT | 961,039,771 | 62,828 | 138x | >10x |
| *fliF-fliRdel* | ONT | 879,718,826 | 60,341 | 126x | >10x |
| *rhbE* | ONT | 425,882,923 | 110,059 | 63x | 62x |
| *rhtA* | ONT | 270,334,017 | 60,801 | 40x | 39x |

### **Table S5.** Bacterial motility modes.

| | Velocity | |
|---|---|---|
| **Motility mode** | **Direction** | **Speed** |
| Swimming | straight | 5 mm/h |
| Trumbling | changes angle | 0 mm/h |
| Swarming | straight | 10 mm/h |
| Entropy-driven swarming | straight | 1 mm/h |
| Twitching | Jerky | 1 mm/h |
| Gliding | jerky | 90 µm/h |
| Sliding | straight | Growth rate x colony size |

The different modes of motility employed by bacteria and their associated speed and direction.

**Table S6.** Comparison of genome sequences of the *fliF-fliR*del mutant to that of the parental *S. meliloti* WT and Sm2011 (GMI11495).

| Replicon | Consensus Position | Type | Length | Consensus | Allele | Count | Coverage | Frequency | Forward/reverse balance | Average quality | Overlapping annotations | Coding region change | Amino acid change |
|---|---|---|---|---|---|---|---|---|---|---|---|---|---|
| chromosome | 1716200 | Insertion | 1 | - | G | 92 | 106 | 86,79245283 | 0,467391304 | 29,07608696 | | | |
| chromosome | 1716251 | Insertion | 7 | - | CGGCCTG | 102 | 106 | 96,22641509 | 0,441176471 | 41,10364146 | | | |
| chromosome | 2059077 | Insertion | 42 | - | GGCGGCGGCGGCGG CGGCGGTGGCGGCG GTGGCGGCGGTGGC | 75 | 82 | 91,46341463 | 0,493333333 | 35,73111111 | CDS: SMc04232 | SMc04232:c.358_359 insGTGGCGGCGGTGG CGGCGGTGGCGGCG GCGGCGGCGGCGGC G | SMc04232: p.Gly119_Asp120insGly GlyGlyGlyGlyGlyGlyGly GlyGlyGlyGlyGlyGly |
| chromosome | 3622706 | SNV | 1 | T | C | 137 | 141 | 97,16312057 | 0,467153285 | 38,87591241 | | | |
| chromosome | 3622726 | MNV | 2 | AA | GG | 137 | 141 | 97,16312057 | 0,481751825 | 40,19708029 | | | |
| chromosome | 3622729 | Deletion | 1 | G | - | 136 | 141 | 96,45390071 | 0,485294118 | 39,91176471 | | | |
| chromosome | 3622746 | SNV | 1 | A | G | 138 | 141 | 97,87234043 | 0,471014493 | 38,21014493 | | | |
| chromosome | 3650478 | SNV | 1 | G | A | 119 | 123 | 96,74796748 | 0,470588235 | 41,6302521 | CDS: SMc02799-rsmG | SMc02799-rsmG:c.151C>T | SMc02799-rsmG: p.His51Tyr |
| pSymB | 726015 | Insertion | 3 | - | CAG | 94 | 103 | 91,26213592 | 0,457446809 | 40,19858156 | CDS: SMb21087-traA2 | SMb21087-traA2:c.33_35dupCA G | SMb21087-traA2: p.Ile11_Arg12insSer |
| pSymB | 726064 | SNV | 1 | G | A | 101 | 103 | 98,05825243 | 0,455445545 | 44,37623762 | CDS: SMb21087-traA2 | SMb21087-traA2:c.82G>A | SMb21087-traA2: p.Ala28Thr |
| pSymB | 726067 | SNV | 1 | A | C | 101 | 103 | 98,05825243 | 0,475247525 | 43,69306931 | CDS: SMb21087-traA2 | SMb21087-traA2:c.85A>C | |
| pSymB | 726090 | SNV | 1 | A | C | 103 | 103 | 100 | 0,466019417 | 45,19417476 | CDS: SMb21087-traA2 | SMb21087-traA2:c.108A>C | |
| pSymB | 726105 | SNV | 1 | C | T | 103 | 103 | 100 | 0,466019417 | 46,44660194 | CDS: SMb21087-traA2 | SMb21087-traA2:c.123C>T | |
| pSymB | 726120 | SNV | 1 | C | G | 103 | 103 | 100 | 0,466019417 | 38,84466019 | CDS: SMb21087-traA2 | SMb21087-traA2:c.138C>G | |
| pSymB | 726123 | SNV | 1 | A | G | 102 | 103 | 99,02912621 | 0,460784314 | 41,81372549 | CDS: SMb21087-traA2 | SMb21087-traA2:c.141A>G | |
| pSymB | 727959 | SNV | 1 | C | T | 107 | 107 | 100 | 0,46728972 | 44,54205607 | CDS: SMb21087-traA2 | SMb21087-traA2:c.1977C>T | |
| pSymB | 729484 | SNV | 1 | T | C | 98 | 109 | 89,90825688 | 0,397959184 | 20,56122449 | CDS: SMb21087-traA2 | SMb21087-traA2:c.3502T>C | |
| pSymB | 729508 | SNV | 1 | G | A | 101 | 110 | 91,81818182 | 0,405940594 | 42,26732673 | CDS: SMb21087-traA2 | SMb21087-traA2:c.3526G>A | SMb21087-traA2:p.Glu1176Lys |
| pSymB | 729543 | MNV | 2 | CA | TG | 110 | 110 | 100 | 0,436363636 | 43,70454545 | CDS: SMb21087-traA2 | SMb21087-traA2:c.3561_3562de linsTG | SMb21087-traA2:p.Thr1188Ala |

| | | | | | | | | | | | | | |
|---|---|---|---|---|---|---|---|---|---|---|---|---|---|
| pSymB | 729548 | SNV | 1 | A | T | 109 | 110 | 99,09090909 | 0,431192661 | 44,40366972 | CDS: SMb21087-traA2 | SMb21087-traA2:c.3566A>T | SMb21087-traA2:p.Tyr1189Phe |
| pSymA | 413296 | SNV | 1 | G | T | 117 | 120 | 97,5 | 0,401709402 | 39,82051282 | CDS: SMa0763 | SMa0763:c.198G>T | SMa0763 (hypothetical protein): p.Leu66Phe |
| pSymA | 568951 | Insertion | 1 | - | A | 119 | 123 | 96,74796748 | 0,462184874 | 41,89915966 | | | |
| pSymA | 711591 | Deletion | 39847 | | | | | | | | fliF to fliR | | |
| pSymA | 911589 | SNV | 1 | A | G | 100 | 102 | 98,03921569 | 0,48 | 38,17 | | | |
| pSymA | 1183827 | SNV | 1 | G | T | 125 | 125 | 100 | 0,416 | 46,68 | CDS: SMa2101 | SMa2101:c.992G>T | SMa2101: p.Gly331Val |

**Legend**

Variation detected both in the *flif-fliR*del mutant and the parental WT when compared to Sm2011 (GMI11495)

Deletion of the *fliF* to *fliR* region (*fliF-fliR*del)

Variation not detected in the parental WT, but in the *flif-fliR*del, *rhbE*, and *rhtA* mutants, indicating that it occurred before mutant constructions

Variation only found in the *flif-fliR*del mutant

**Table S7.** Comparison of genome sequences of the *rhbE* mutant to that of the parental *S. meliloti* WT and Sm2011 (GMI11495).

| Mapping | Con-sensus Position | Type | Length | Consensus | Allele | Count | Cove-rage | Frequency | Forward/reverse balance | Average quality | Overlapping annotations | Coding region change | Amino acid change |
|---|---|---|---|---|---|---|---|---|---|---|---|---|---|
| chromo-some | 1716200 | Insertion | 1 | - | G | 48 | 59 | 81,3559322 | 0,35416667 | 28,2083333 | | | |
| chromo-some | 1716251 | Insertion | 7 | - | CGGCCTG | 54 | 59 | 91,5254237 | 0,42592593 | 38,8624339 | | | |
| chromo-some | 2059077 | Insertion | 42 | - | GGCGGCGGCGGCGGCGGCGGTGGCGGCGGTGGCGGCGGTGGC | 32 | 38 | 84,2105263 | 0,46875 | 39,4501488 | CDS: SMc04232 | SMc04232:c.358_359insGTGGCGGCGGTGGCGGCGGTGGCGGCGGCGGCGGCGGCGGCG | SMc04232:p.Gly119_Asp120insGlyGlyGlyGlyGlyGlyGlyGlyGlyGlyGlyGlyGlyGly |
| chromo-some | 3622706 | SNV | 1 | T | C | 87 | 88 | 98,8636364 | 0,48275862 | 34,6206897 | | | |
| chromo-some | 3622726 | MNV | 2 | AA | GG | 89 | 89 | 100 | 0,48314607 | 37,3764045 | | | |
| chromo-some | 3622729 | Deletion | 1 | G | - | 88 | 89 | 98,8764045 | 0,48863636 | 36,5909091 | | | |
| chromo-some | 3622746 | SNV | 1 | A | G | 90 | 90 | 100 | 0,48888889 | 35,3222222 | | | |
| chromo-some | 3650478 | SNV | 1 | G | A | 88 | 92 | 95,6521739 | 0,47727273 | 42,1590909 | CDS: SMc02799-rsmG | SMc02799-rsmG:c.151C>T | SMc02799-rsmG:p.His51Tyr |
| pSymB | 726015 | Insertion | 3 | - | CAG | 51 | 59 | 86,440678 | 0,47058824 | 37,4313725 | CDS: SMb21087-traA2 | SMb21087-traA2:c.33_35dupCAG | SMb21087-traA2:p.Ile11_Arg12insSer |
| pSymB | 726064 | SNV | 1 | G | A | 59 | 59 | 100 | 0,44067797 | 40,1186441 | CDS: SMb21087-traA2 | SMb21087-traA2:c.82G>A | SMb21087-traA2:p.Ala28Thr |
| pSymB | 726067 | SNV | 1 | A | C | 59 | 59 | 100 | 0,44067797 | 38,9661017 | CDS: SMb21087-traA2 | SMb21087-traA2:c.85A>C | |
| pSymB | 726090 | SNV | 1 | A | C | 59 | 60 | 98,3333333 | 0,45762712 | 42,9830508 | CDS: SMb21087-traA2 | SMb21087-traA2:c.108A>C | |
| pSymB | 726105 | SNV | 1 | C | T | 62 | 62 | 100 | 0,4516129 | 41,8548387 | CDS: SMb21087-traA2 | SMb21087-traA2:c.123C>T | |

| | | | | | | | | | | | | | |
|---|---|---|---|---|---|---|---|---|---|---|---|---|---|
| pSymB | 726120 | SNV | 1 | C | G | 62 | 62 | 100 | 0,4516129 | 35,1612903 | CDS: SMb21087-traA2 | SMb21087-traA2:c.138C>G | |
| pSymB | 726123 | SNV | 1 | A | G | 61 | 62 | 98,3870968 | 0,45901639 | 37,8360656 | CDS: SMb21087-traA2 | SMb21087-traA2:c.141A>G | |
| pSymB | 727959 | SNV | 1 | C | T | 55 | 57 | 96,4912281 | 0,45454545 | 38,4909091 | CDS: SMb21087-traA2 | SMb21087-traA2:c.1977C>T | |
| pSymB | 729484 | SNV | 1 | T | C | 57 | 62 | 91,9354839 | 0,40350877 | 18,754386 | CDS: SMb21087-traA2 | SMb21087-traA2:c.3502T>C | |
| pSymB | 729508 | SNV | 1 | G | A | 60 | 62 | 96,7741935 | 0,41666667 | 39,75 | CDS: SMb21087-traA2 | SMb21087-traA2:c.3526G>A | SMb21087-traA2:p.Glu1176Lys |
| pSymB | 729543 | MNV | 2 | CA | TG | 62 | 62 | 100 | 0,43548387 | 41,9919355 | CDS: SMb21087-traA2 | SMb21087-traA2:c.3561_3562delinsTG | SMb21087-traA2:p.Thr1188Ala |
| pSymB | 729548 | SNV | 1 | A | T | 61 | 62 | 98,3870968 | 0,42622951 | 42,0655738 | CDS: SMb21087-traA2 | SMb21087-traA2:c.3566A>T | SMb21087-traA2:p.Tyr1189Phe |
| pSymB | 838025 | Deletion | 1 | G | - | 45 | 53 | 84,9056604 | 0,44444444 | **13,4** | CDS: SMb21264 | SMb21264:c.881delG | SMb21264 (putative glycosyl transferase): p.Gly294fs |
| pSymA | 568951 | Insertion | 1 | - | A | 56 | 59 | 94,9152542 | 0,5 | 38,7321429 | | | |
| pSymA | 616106 | Deletion | 1 | T | - | 70 | 71 | 98,5915493 | 0,47142857 | 34,1571429 | | | |
| pSymA | 911589 | SNV | 1 | A | G | 47 | 50 | 94 | 0,40425532 | 37,7446809 | | | |
| pSymA | 1183827 | SNV | 1 | G | T | 60 | 60 | 100 | 0,45 | 40,8666667 | CDS: SMa2101 | SMa2101:c.992G>T | SMa2101:p.Gly331Val |
| pSymA | 1302927 | Deletion | 2 | TC | - | 61 | 77 | 79,2207792 | 0,49180328 | 31,4918033 | | | |
| pSymA | 1309739 | Deletion | 1354 | ATCATGACGGATTTCGATCTGGCAGGGATCGGCATCGGGCCGTTCAACCTTGGCCTTGCCGCCCTTCTCTCTTCCCACGAAAATCTCTCGAACGTGTTTCTCGAGCGCAAACCGGCGTTCCGGTGGCACGAAGGGCTCATTCTTCCCGGGACGACGCTGCAGGTTCCCTTCATGGCGGATCTCGTCACCATGGCGGACCCGACGCATCGTTTGAGCTTCCTCAACTACCTTGCTGTGCACGATCGCCTCTACAAGTTCTACTTCTACGAAAACTTCATGATCCCGCGGCAGGAATACGATCACTATTGCCGCTGGGCCTCACAGCAACTTTCGGCGTGCCGCTTCG | - | 48 | 50 | 96 | 0,5 | 34,3125 | CDS: SMa2406-rhbD, CDS: SMa2408-rhbE | SMa2406-rhbD:c.591_*1340del; SMa2408-rhbE:c.4_1357del | SMa2406-rhbD:p.*197fs; SMa2408-rhbE:p.Ile2_Ser448del |

| | | | | |
|---|---|---|---|---|
| | | | | GCGAGGAAGTGGTCGACGTCGCCCAT<br>GAGTCGGCATCCGACAGCTTCATTGTC<br>GAAAGCAGGTCCGCCTCGGGCGGAAA<br>GCAGCAATATCGAAGCCGTAATATTG<br>CGATCGGCGTCGGCACGGCTCCTTTCC<br>TGCCGAAGTGGGCGCAGATCAAGACG<br>TTGGCACCGCTCATGCATTCCAGCGAA<br>TTCGGGAGGCGGCGTCTGGAGCTTTC<br>GAAACGACGCCGTGTAACAGTCATCG<br>GCTCAGGGCAAAGCGCTGCCGAATGC<br>GTTCTTGCGCTTTTGAACGATCTGACG<br>CCGGAGATGGTAGCAGCCGGCGCTTC<br>GATCCAGTGGATCACCCGATCGGCGG<br>GGTTCTTTCCGATGGAGTATTCGAAGC<br>TGGGCCTCGAATACTTCACGCCTGACT<br>ACATGCGCCACTTCCATCGAATCGCGC<br>CCGTTAGACGCCGCGAGATCGTTGCA<br>GACCAGGGCCTGCTCTACAAGGGCAT<br>CAGCTTTTCGACGATCGGTGAGATCTT<br>CGATCTCATGTATGAACGTTCGGTGG<br>GCGGACGCGATCCAGGCCTAGCACTC<br>TTTTCCAATTGTGCGGTCGAAACGCTG<br>GAGAGCGCAGGGGGATCGGGGTCGT<br>TTCGCATCGGCATCAACCACAACCATC<br>TGGATGAGAAGGCAACGGTCGAGACC<br>GACGCGATTGTGGCCGCAACGGGGTA<br>CCGGCACGCGTGGCCGGAATGGCTGG<br>GTTCGCTGAAGGGCAGCGTCCTCGAT<br>ACCTGCGAGTGGGGCGACCTCGTCGT<br>CGGCGGGGATTTCCGTGCACGCCGCA<br>GCGACGGCGGCAAGGGCCATGTGTTC<br>GTCCAGAATGCCGAAACCTTCCACCAC<br>GGCGTCGGCGCGCCGGATCTTGGCCT<br>CGGCGCCTTTCGCAATGCCGTCATCGT<br>CAATCAGCTTCTCGGTCGCGAGCATTA<br>CCGCGTCAATGCGTCGGCATCGTTCCA<br>GAAGTTCGGTCTGCCCTCCAGCCAGA<br>CCGCTCCGTCCTCGATTTCA |

**Legend**

Variation detected both in the *rhbE* mutant and the parental WT when compared to Sm2011 (GMI11495)

Deletion of *rhbE* (includes deletion of the third base of the stop codon of the upstream *rhbD* gene; a stop codon is reconstituted by fusion to the 3'-end of deletion resulting in a functional *rhbD* cds)

Variation not detected in the parental WT, but in the *flif-fliR*del, *rhbE*, and *rhtA* mutants, indicating that it occurred before mutant constructions

Variation only found in the *rhbE* mutant

**Indicates low sequencing quality area (GC rich)**

**Table S8.** Comparison of genome sequences of the *rhtA* mutant to that of the parental *S. meliloti* WT and Sm2011 (GMI11495).

| Map-ping | Con-sensus Position | Type | Length | Consensus | Allele | Count | Cove-rage | Frequency | Forward/reverse balance | Average quality | Overlapping annotations | Coding region change | Amino acid change |
|---|---|---|---|---|---|---|---|---|---|---|---|---|---|
| chromo-some | 1716200 | Insertion | 1 | - | G | 38 | 43 | 88,372093 | 0,5 | 26,2105263 | | | |
| chromo-some | 1716251 | Insertion | 7 | - | CGGCCTG | 40 | 43 | 93,0232558 | 0,475 | 39,7428571 | | | |
| chromo-some | 2059077 | Insertion | 42 | - | GGCGGCGGCGGCGGCGGCGGTGGCGGCGGTGGCGGCGGTGGC | 15 | 19 | 78,9473684 | 0,266666667 | 37,8428571 | CDS: SMc04232 | SMc04232:c.358_359insGTGGCGGCGGTGGCGGCGGTGGCGGCGGCGGCGGCGGCGGCG | SMc04232:p.Gly119_Asp120insGlyGlyGlyGlyGlyGlyGlyGlyGlyGlyGlyGlyGlyGly |
| chromo-some | 3622706 | SNV | 1 | T | C | 46 | 48 | 95,8333333 | 0,47826087 | 35,4782609 | | | |
| chromo-some | 3622726 | MNV | 2 | AA | GG | 47 | 47 | 100 | 0,489361702 | 38,6808511 | | | |
| chromo-some | 3622729 | Deletion | 1 | G | - | 47 | 47 | 100 | 0,489361702 | 37,2765957 | | | |
| chromo-some | 3622746 | SNV | 1 | A | G | 46 | 47 | 97,8723404 | 0,47826087 | 36,0869565 | | | |
| chromo-some | 3650478 | SNV | 1 | G | A | 40 | 44 | 90,9090909 | 0,45 | 42,75 | CDS: SMc02799-rsmG | SMc02799-rsmG:c.151C>T | SMc02799-rsmG:p.His51Tyr |
| pSymB | 726015 | Insertion | 3 | - | CAG | 31 | 35 | 88,5714286 | 0,419354839 | 38,0322581 | CDS: SMb21087-traA2 | SMb21087-traA2:c.33_35dupCAG | SMb21087-traA2:p.Ile11_Arg12insSer |
| pSymB | 726064 | SNV | 1 | G | A | 34 | 34 | 100 | 0,441176471 | 41,7941176 | CDS: SMb21087-traA2 | SMb21087-traA2:c.82G>A | SMb21087-traA2:p.Ala28Thr |
| pSymB | 726067 | SNV | 1 | A | C | 34 | 34 | 100 | 0,441176471 | 40,1764706 | CDS: SMb21087-traA2 | SMb21087-traA2:c.85A>C | |
| pSymB | 726090 | SNV | 1 | A | C | 32 | 32 | 100 | 0,4375 | 44,25 | CDS: SMb21087-traA2 | SMb21087-traA2:c.108A>C | |

| | | | | | | | | | | | | | |
|---|---|---|---|---|---|---|---|---|---|---|---|---|---|
| pSymB | 726105 | SNV | 1 | C | T | 31 | 32 | 96,875 | 0,419354839 | 44,6451613 | CDS: SMb21087-traA2 | SMb21087-traA2:c.123C>T | |
| pSymB | 726120 | SNV | 1 | C | G | 31 | 31 | 100 | 0,451612903 | 36,3548387 | CDS: SMb21087-traA2 | SMb21087-traA2:c.138C>G | |
| pSymB | 726123 | SNV | 1 | A | G | 31 | 31 | 100 | 0,451612903 | 38,5806452 | CDS: SMb21087-traA2 | SMb21087-traA2:c.141A>G | |
| pSymB | 727959 | SNV | 1 | C | T | 37 | 38 | 97,3684211 | 0,486486486 | 39,7567568 | CDS: SMb21087-traA2 | SMb21087-traA2:c.1977C>T | |
| pSymB | 729484 | SNV | 1 | T | C | 30 | 31 | 96,7741935 | 0,366666667 | 18,2333333 | CDS: SMb21087-traA2 | SMb21087-traA2:c.3502T>C | |
| pSymB | 729508 | SNV | 1 | G | A | 30 | 31 | 96,7741935 | 0,366666667 | 37,9 | CDS: SMb21087-traA2 | SMb21087-traA2:c.3526G>A | SMb21087-traA2:p.Glu1176Lys |
| pSymB | 729543 | MNV | 2 | CA | TG | 31 | 31 | 100 | 0,387096774 | 43,8548387 | CDS: SMb21087-traA2 | SMb21087-traA2:c.3561_3562delinsTG | SMb21087-traA2:p.Thr1188Ala |
| pSymB | 729548 | SNV | 1 | A | T | 31 | 31 | 100 | 0,387096774 | 40,8387097 | CDS: SMb21087-traA2 | SMb21087-traA2:c.3566A>T | SMb21087-traA2:p.Tyr1189Phe |
| pSymA | 568951 | Insertion | 1 | - | A | 42 | 44 | 95,4545455 | 0,476190476 | 38,3809524 | | | |
| pSymA | 911589 | SNV | 1 | A | G | 30 | 30 | 100 | 0,466666667 | 34,7 | | | |
| pSymA | 1183827 | SNV | 1 | G | T | 34 | 34 | 100 | 0,5 | 42,7941176 | CDS: SMa2101 | SMa2101:c.992G>T | SMa2101:p.Gly331Val |

| | | | | | | | | | | | | | |
|---|---|---|---|---|---|---|---|---|---|---|---|---|---|
| pSymA | 1314087 | Deletion | 2235 | CAACAATGAGAATGGCGGCATAAGCTTTTGCGTCTTCGTCGTCGTAATTGGATTTGGCACGGGTGCTGTTGCGCAGGAGCCAGCGAATCAATCCGAAGCTGTGACGAGCCTGGAAGAAATCGTAGTCACCGGCGGACGATCCGCTCAGCAGATTTCTGAAATCGCGCGCACCATTTACGTCGTCGATTCGGATCAGATCCAGGCCGAGGCCCGGTCGGGCAAGACGCTGCAGCAGATTCTGGGCGAAACCATTCCGAGCTTCGACCCCGCCAGCGACGGAGCGCGTACGTCATTCGGCCAGAACCTGCGTGGCCGGCCACCACTGATCCTTGTCGACGGTGTCTCGATGAACTCGGCGCGCTCGCTGAGCCGACAATTCGATGCCATCGATCCATTCAACATCGAACGGGTGGAGGTCCTGTCTGGGGCAACTGCAATCTACGGCGGGAACGCAACCGGCGGCATCATCAACATCATCACGAAGAAGGGCAAGGACGCGGAACCGGGCCTGCATGCGGAAGTGACCGGCGGCATGGGCAGCGGCTTTGCCGGCAGCCAGGACTTCGACCGCAATGCGGCGGGTGCGGTCACCTATAACAGCGAAAACTGGGATGCCCGCCTTTCGATCGCCGGCAACAGGACCGGCGCCTTTTATGACGGCAGCGGCACGCTGCTGATCCCCGACATCACGCAGACGTCCACCGCATTCAACGAGCGCATCGACCTGATGGGGTCTATCGGCTACCAGATCGACGACGACCGTCGCGTCGAATTTTCCGGGCAATATTTCGACAGCAAGCAGGACTCGGATTACGGGCTTTATTACGGGCCCTTCTTTGCAGCGCTCGCCGACCCGAGCCTGTTTGAAACCCGTTCCGGATACGAGTCCGACTTCAATCCGCAGACACGCCGCTCGATGTTGAACGTCACCTATACCGATAATGACGTTTTCGGTCAGCAGCTATTGCTTCAGGGATCTTACCGGACTGAGCGGATAAAGTTCCATCCCTTCCCTGCTTCGGGCAATAGCGAAACGGGTCCTTACTTTTACGGCAGTTCGCAGGACACAGACTATTACGGCATCAGGGCTGCCTTGGTTGCGGAACCCACCGACGCGCTGAAAATCACCTATGGCATCGATGCGGATATGGATTCCTTCACGGCGCGTCAGAACATCTTCGACATGGTGGCAGCCGGGCAATCCGGGGGGC | - | 33 | 36 | 91,6666667 | 0,454545455 | 31,5151515 | CDS: SMa2414-rhtA | SMa2414-rhtA:c.6_2240del | SMa2414-rhtA:p.Asn3_*747del |

| | | | | | | | | | | | | | |
|---|---|---|---|---|---|---|---|---|---|---|---|---|---|
| | | | | TCGATTTCAATACGATCGGCAAGACC<br>GGTCTTTATCCATCCATCGATGTCTCC<br>ACCGTTGCCGGCTTTGCTGAGGCCTCG<br>TACGAGGCTACCGACAGGCTCACTCT<br>GAACGGCGGGGTGCGCTATCAGTTCG<br>TCAACACTGAGGTATCCGACTTCATCG<br>GAGCGGCTCAGCAGGTCGCCATCCTG<br>CAGGGCAGAGCAACGTCTGCCGACAC<br>CATACCTGGCGGCGAAGTCAATTACG<br>ACGCCGCTTTGTTCAGTGCAGGCGCA<br>ACCTACCAGTTGACGAATACGCAGCA<br>GGTCTACGCCAATTTCAGCCAGGGCTT<br>CGAACTACCAGATCCGGCCAAGTACT<br>ATGGAATTGGCAACTACTCCTTCTCCG<br>GTGGACACTACACGCTTGTCAATAGC<br>GTGAACGTGGGCGATTCCGCGCTCGA<br>GGCGATCAAGACCAACTCATTCGAAA<br>TCGGCTATCGTCTCGATGACGGCACCT<br>TCAATCTTGAAACGGCGGCCTACTACT<br>CGCTCTCCGATCGTTCCATCAATCTCA<br>ATCGCTCATCGCTTGCCGTTGAGATCA<br>TCGATCGGGAGAGACGCGTCTATGGC<br>ATCGAAGGAAAGGCGGGCGTAAAGC<br>TCGATCATGGCTTCGATGTTGGAGTTC<br>TGGGGCACTGGGTCAGAACCGAGGTC<br>AAGGGAGCCGACGGTTGGGAAAAGG<br>ACTCCGTCGGCAGCGCGAGCGTCTCC<br>AAACTCGGCGGCTATGTCGGCTGGAC<br>CAATGACGCTCTCAGCCTGAGGTTTTC<br>CGGTCAGCACATCTTCGAACTCACCGA<br>CGCCCAGAATTTCACGATCGACGATTA<br>TACGTTGTTTGATCTAACCGGCGGCTA<br>CAGGTTCGAGAATACCGACACGACGC<br>TGAATTTCGGTATCCATAACGTCTTCG<br>ATACTGATTACACCACCGTCTGGGGCT<br>CTCGCGCCAAAGCCCTTTACGGCGGC<br>CTAGCCGACGAGTCGGTGTTCGACTA<br>CAAGGGCCGCGGCCGCACGTTCGCGG<br>TCTCGCTGACAAAGGTTTTTTA | | | | | | | | | |
| pSymA | 1316324 | Replace-ment | 2 | G | CC | 33 | 36 | 91,6666667 | 0,454545455 | 36,4848485 | CDS: SMa2339 (located downstream of rhtA) | SMa2339:c.3delinsCC (putative change of the ATG start codon to ACC) | SMa2339 (siderophore biosynthesis protein): p.Met1? |

**Legend**

Variation detected both in the *rhtA* mutant and the parental WT when compared to Sm2011 (GMI11495)

Deletion of *rhtA*

Variation not detected in the parental WT, but in the *flif-fliR*del, *rhbE*, and *rhtA* mutants, indicating that it occurred before mutant constructions

Variation only found in the *rhtA* mutant

**Supplemental Figures:**

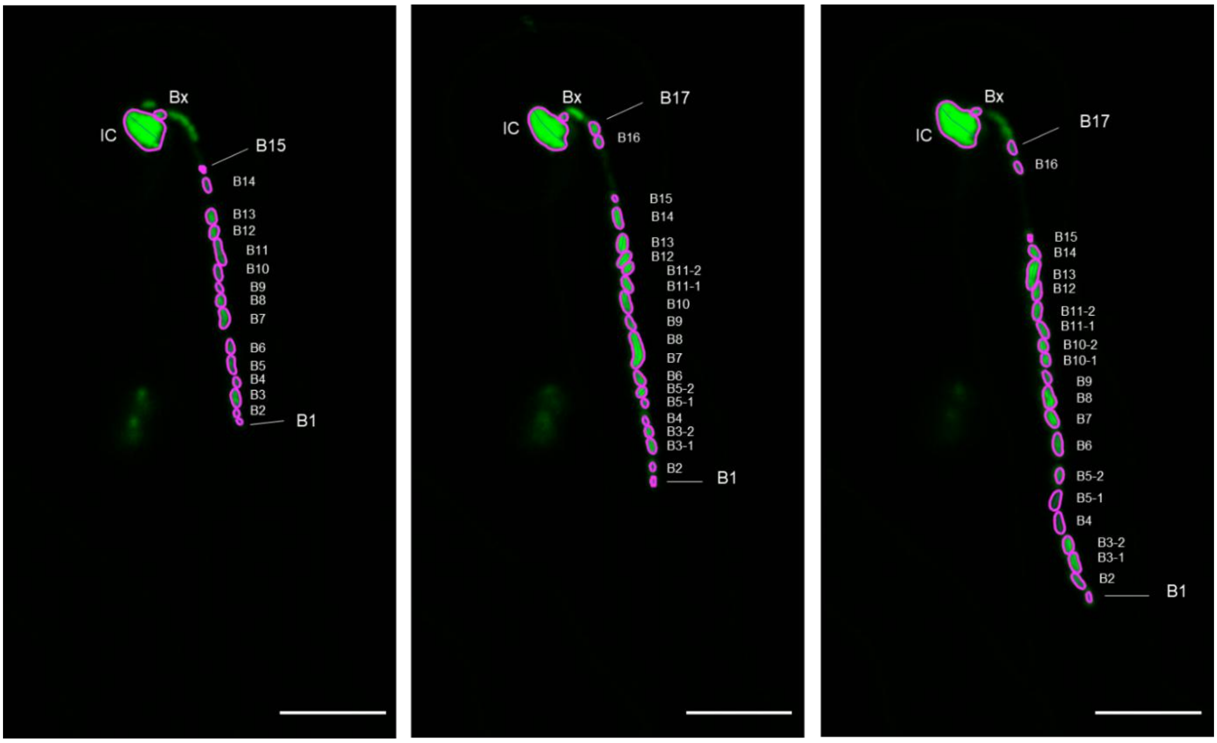

### **Fig. S1.** Tracking bacteria within ITs.

Regions of interest (ROIs) corresponding to individual bacteria (magenta) were created using the MicrobeJ plugin of ImageJ. Each rhizobial cell in the file was numbered starting from the IT tip (B1, B2, etc), at time point 1. Bacteria that had divided during a time interval (B3, B5, B10, B11) were named after the parent cell. The ROIs corresponding to the infection chamber (IC) and a bacterium close to it (Bx) were used as spatial references. When individual bacteria could not be distinguished (see B7 and B8 at time point 2) or a bacterium divided over the next time interval (see B3, for example), their speed was not measured (and is thus missing in Fig.1b). Spatial coordinates of each ROI were determined and exported using MicrobeJ.

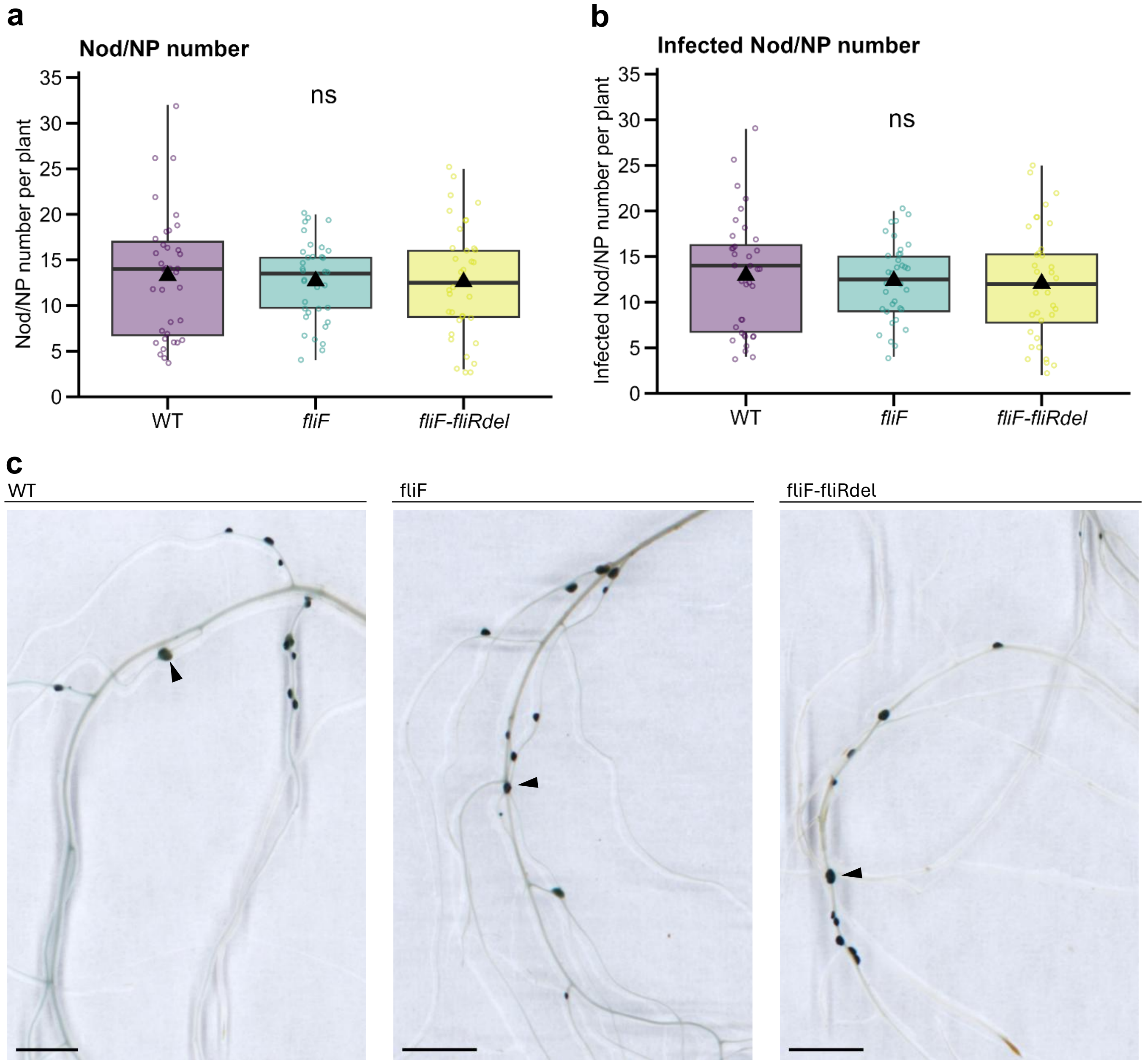

**Fig. S2.** *S. meliloti fliF* and *fliF-fliRdel* flagella motility mutants form nodules fully colonized by rhizobia in *Medicago truncatula* in different growth conditions.

*M. truncatula* plants grown in inert attapulgite substrate were inoculated with lacZ-expressing *S. meliloti* WT or *fliF* and *fliF-fliRdel* mutant strains. Number of nodules (Nod) and nodule primordia (NP) per plant (**a**) and of infected Nod/NP per plant (**b**) were quantified in nodulated roots using a binocular magnifying glass at 7 dpi (WT n = 36; *fliF* n = 36; *fliF-fliRdel* n = 36). Box plots (**a-b**) show the distribution of values (circles) from 3 independent experiments. First and third quartiles (horizontal box edges), minimum and maximum (whisker tips), median (centerline), mean (solid black triangle), and outliers (crosses) are shown. ns indicate no statistical difference relative to WT ($p$ = 0,894 in **a**, $p$ = 0,799 in **b;** one-way ANOVA). (**c**) Representative images of LacZ-stained roots of *M. truncatula* at 7 dpi with WT and mutant *S. meliloti* strains. Fully-infected nodules (arrowheads) are observed in both WT and mutant-inoculated plants. Scale bars in (**c**) = 3 mm.

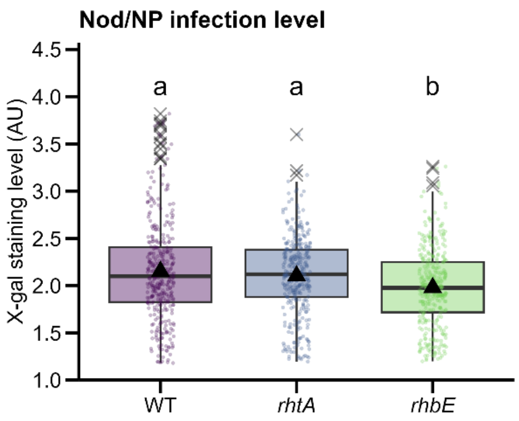

**Fig. S3.** Mutation in *rhtA* Rhizobactin 1021 transporter does not affect early rhizobia nodule colonization.

*M. truncatula* A17 grown in vermiculite:sand pots were inoculated with *S. meliloti* WT, *rhtA and rhbE* mutant strains and quantification of Nod/NP infection level (X-gal staining intensity) was performed 7 dpi in individual nodules (WT n = 352; *rhtA* n = 342; *rhbE* n = 357). Box plots show the distribution of values (circles) from 2 independent experiments. First and third quartiles (horizontal box edges), minimum and maximum (whisker tips), median (centerline), mean (solid black triangle), and outliers (crosses) are shown. Letters indicate statistically significant differences between groups ($p$ = 6.276046e-06**,** Kruskal-Wallis $\alpha$ = 5%).
